# Recombination rate variation shapes barriers to introgression across butterfly genomes

**DOI:** 10.1101/297531

**Authors:** Simon H. Martin, John W. Davey, Camilo Salazar, Chris D. Jiggins

**Affiliations:** Department of Zoology, University of Cambridge, Cambridge CB2 3EJ, United Kingdom; Department of Biology, University of York, YO10 5DD, United Kingdom; Biology Program, Faculty of Natural Sciences and Mathematics, Universidad del Rosario, Carrera. 24 No. 63C-69, Bogota, D.C. 111221, Colombia

## Abstract

Hybridisation and introgression can dramatically alter the relationships among groups of species, leading to phylogenetic discordance across the genome and between populations. Introgression can also erode species differences over time, but selection against introgression at certain loci acts to maintain post-mating species barriers. Theory predicts that species barriers made up of many loci throughout the genome should lead to a broad correlation between introgression and recombination rate, which determines the extent to which selection on deleterious foreign alleles will affect neutral alleles at physically linked loci. Here we describe the variation in genealogical relationships across the genome among three species of *Heliconius* butterflies: *H. melpomene*, *H. cydno* and *H. timareta*, using whole genomes of 92 individuals, and ask whether this variation can be explained by heterogeneous barriers to introgression. We find that species relationships vary predictably at the chromosomal scale. By quantifying recombination rate and admixture proportions, we then show that rates of introgression are predicted by variation in recombination rate. This implies that species barriers are highly polygenic, with selection acting against introgressed alleles across most of the genome. In addition, long chromosomes, which have lower recombination rates, produce stronger barriers on average than short chromosomes. Finally, we find a consistent difference between two species pairs on either side of the Andes, which suggests differences in the architecture of the species barriers. Our findings illustrate how the combined effects of hybridisation, recombination and natural selection, acting at multitudes of loci over long periods, can dramatically sculpt the phylogenetic relationships among species.

## INTRODUCTION

The genealogical relationships among closely-related species can be complex, varying across the genome and among individuals. This heterogeneity is most prevalent in the presence of introgressive hybridisation, which can alter species’ relationships in certain parts of the genome. Genome-scale studies have revealed that particular genomic regions such as sex chromosomes and chromosomal inversions can have distinct phylogenetic histories [1–3], indicating heterogeneity in introgression across the genome. Indeed, the establishment of barriers to introgression in certain parts of the genome through selection against hybrids or admixed individuals is a key part of the speciation process [4–7]. Selection against foreign genetic variation can be driven by both extrinsic factors, such as adaptation to distinct environments, and intrinsic factors, such as genetic incompatibilities [6,8]. The heterogeneous landscape of species relationships therefore carries information about the ‘barrier loci’ that contribute to the origin and maintenance of species.

The barriers between closely-related subspecies or ecotypes that interbreed frequently are often restricted to just a few loci that contribute to local adaptation, resulting in narrow ‘islands’ of genetic differentiation between populations [1,9–11]. Recently, it has become evident that patterns of genomic differentiation between more strongly isolated species are often complex and result from an interaction of genomic processes including localised selective sweeps and background selection within species that reduce variation at linked sites [12–14]. This is particularly the case for conventional measures of genetic differentiation, such as *F*_ST_, which provide a poor proxy for the strength of a local barrier. However, it is possible to largely avoid the confounding effects of positive and background selection by directly estimating the effective migration rate and how it varies across the genome, either using summary statistics that are less prone to artefacts [15], or through model-based inference [16]. Provided there has been sufficient migration between the species, regions of the genome where admixture is reduced can be inferred to have experienced selection against foreign genetic variation.

Once beyond the very earliest stages of divergence, post-mating species barriers could involve many loci (‘polygenic’ barriers) [17]. If species barriers are highly polygenic and each locus has only a weak effect on fitness, their individual localised effects on levels of admixture might be difficult to detect, analogous to the difficulties in studying polygenic adaptation more generally [18– 20]. While it may not be possible to identify all barrier loci in such a situation, we can test hypotheses about the architecture of barriers by studying genome-wide patterns of admixture. In particular, barriers made up of many loci of small effect are expected to be weaker where recombination rates are higher. Foreign chromosomes that enter a population through hybridisation and backcrossing will be more rapidly broken down over subsequent generations in regions with higher recombination rates. This creates more opportunities for neutral (or mutually beneficial) foreign alleles to be separated from detrimental foreign alleles at other loci, and thus avoid being removed by selection [16,21–23]. A correlation between the recombination rate and the inferred rate of effective migration has been observed between subspecies of house mice [24], subspecies of *Mimulus* monkeyflowers [16] and even between humans and Neanderthals [25], suggesting that loci experiencing selection against introgression among close relatives can be widespread in the genome. To date, it has not been investigated whether such widespread barrier loci could also explain large-scale heterogeneity in phylogenetic relationships across the genome.

We explored species relationships and barriers to introgression among species of *Heliconius* butterflies. Many *Heliconius* species are divided into geographically distinct ‘races’ with distinct warning patterns, which signal their distastefulness to local predators. Selection favouring locally recognised warning patterns maintains narrow islands of divergence at a few wing patterning loci between otherwise genetically similar races [1,9,26]. However, there are also more strongly differentiated pairs of sympatric species that hybridise rarely and have strong post-zygotic barriers, leading to higher genome-wide genetic differentiation [1]. We studied three such species: *Heliconius melpomene* (‘*mel’*), *Heliconius cydno* (‘*cyd’*) and *Heliconius timareta* (‘*tim’*), which form at least two independent zones of sympatry separated by the Andes mountains. While *mel* is found throughout much of South and Central America, *cyd* is largely restricted to the west of the Andes and the inter-Andean valleys, where it overlaps with the western populations of *mel*, whereas *tim* occurs only on the eastern slopes of the Andes, where it co-occurs with the eastern populations of *mel*. In addition to strong assortative mating based on chemical cues and wing patterns in the case of *cyd* and *mel* [27–30], and mainly chemical cues between *mel* and *tim* [29,31,32], both species pairs show ecological differences as well as partial hybrid sterility [29,30,33–37] (and see [29] for a review). Nevertheless, previous studies have revealed surprisingly pervasive admixture between these species in sympatry, most likely explained by a low rate of ongoing hybridisation over an extended period of time [1,38,39]. There is also considerable heterogeneity in the relationships among these populations across the genome [1]. Adaptive introgression in *Heliconius* is well documented. Mimicry between sympatric races of *mel* and *tim* has been facilitated by exchange of multiple wing patterning alleles [40,41], and at least one case of introgression between *mel* and *cyd* has allowed the latter to mimic other unpalatable species [42]. However, the extent to which introgression among these species might be selected against remains unclear.

Using 92 whole genome sequences, we asked whether the heterogeneous relationships observed among these species reflect the influence of polygenic barriers to introgression that vary in their strength across the genome. Then, taking advantage of high-resolution linkage maps for these species [43], we show that admixture is correlated with recombination rate, consistent with polygenic species barriers leading to widespread selection against introgression. This selection also explains broader variation in admixture at the chromosomal scale. Overall, our results highlight the pervasive role of natural selection in shaping the ancestry of hybridising species.

## RESULTS

### Population Structure

We analysed whole genome sequence data from 90 butterflies representing nine populations of *H. melpomene* (‘*mel’*, 50 samples from five populations or ‘races’), *H. cydno* (‘*cyd’*, 20 samples from two races) and *H. timareta* (‘*tim’*, 20 samples from two races), along with two samples from an outgroup species *Heliconius numata* (‘*num’*) (Table S1). Our sampling included four regions of sympatry: two on the west of the Andes where *cyd* co-occurs with *mel-W*, and two on the eastern slopes of the Andes where *tim* co-occurs with *mel-E*, as well as an allopatric population, *mel-G*, from French Guiana (Figure 1A). Principal components analysis (PCA) based on whole-genome SNP data shows clear distinctions between the three species, and also between *mel-W*, *mel-E* and *mel-G* (Figure 1B). By contrast, pairs of races of the same species from the same broad geographic area (i.e. ‘West’, ‘East’ and ‘Guiana’) are not clearly distinct in the PCA, indicating virtually panmictic populations in each species in each area, despite variation at a few wing patterning loci, as shown previously [44,45]. These results therefore highlight the contrast between the clear barriers that exist between sympatric species, even in sympatry, and the continuity that exists within species, with the Andes mountains and wide Amazon basin presenting the only major sources of discontinuity among sampled populations of the same species [44,45].

**Figure 1.**
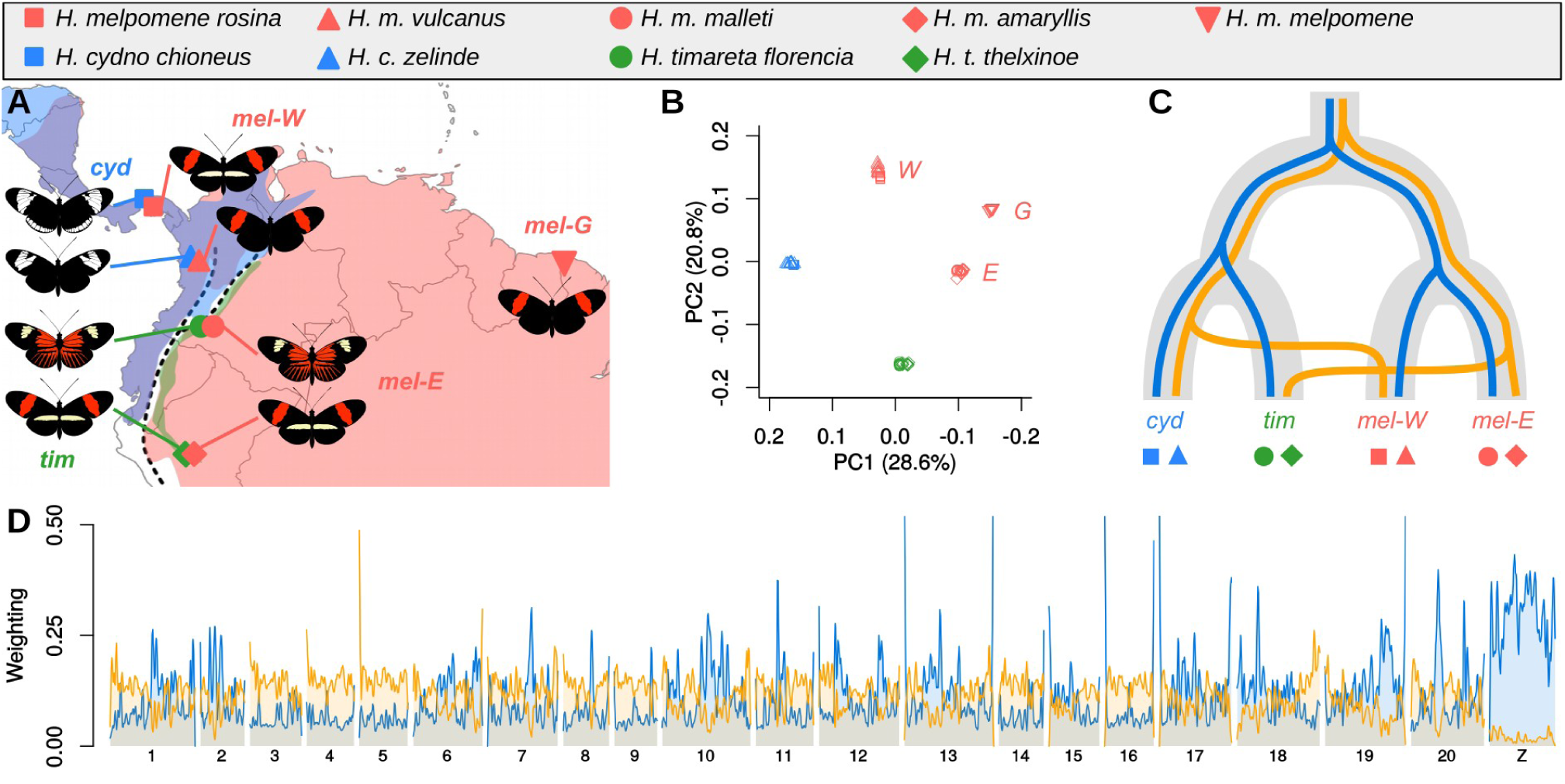
The three species are clearly distinct but their relationships vary across the genome. **A.** Sampling locations of the nine races from three species included in this study. Species ranges for *H. melpomene*, *H. cydno* and *H. timareta* are indicated by red, blue and green shading, respectively. The dashed line indicates the central part of the Andean mountains **B.** Principal components 1 and 2 differentiate both *cyd* and *tim* from three *mel* populations, but do not separate the two sampled races of each species on either side of the Andes. **C.** The two topologies with the highest weightings, the species topology (‘T3’, blue) and geography topology (‘T6’, yellow), shown here as lineages within the hypothesized population branching history. While the geography can arise through both introgression and lineage sorting effects, here it is represented as arising through recent introgression, with the direction (i.e. from *mel-E* into *tim* and from *cyd* into *mel-W*) representing the inferred prevailing direction of introgression, as described below. **D.** Weightings for the ‘species’ and ‘geography’ topologies plotted across the 21 chromosomes and smoothed as a locally-weighted average (loess span = 1 Mb). See Figure S4 for a detailed plot without smoothing.

### Topology weighting reveals both introgression and incomplete linage sorting

We explored species relationships across the genome using *Twisst* [46], which quantifies the frequency (or ‘weighting’) of alternative topological relationships among all sampled individuals in narrow windows of 50 single nucleotide polymorphisms (SNPs) each. Consistent with previous results, topology weighting indicates that both large-scale introgression and stochasticity in lineage sorting have shaped the relationships among these species. All 15 possible rooted topologies that describe the relationship between *cyd*, *tim*, *mel-W* and *mel-E* (rooted with *H. numata* as the outgroup) (Figure S1) have non-zero weightings (Figure S2). Despite the strong clustering of distinct species in the PCA, less than 0.5% of windows have completely-sorted genealogies (i.e. in which all groups cluster according to a single topology, resulting in a weighting of one) (Figure S2). This low level of lineage sorting is not surprising given the large effective population sizes (> 2 million individuals [45]), fairly recent divergence times, and large sample sizes used. The two most common topologies are 3 and 6, which differ entirely in the relationships among the ingroup taxa (Figure 1C). Topology 3 matches the expected species branching order, in which *cyd* and *tim* are sister species and *mel*-W groups with *mel*-E ((*cyd*, *tim*), (*mel-W*, *mel-E*)) [44,47]. We refer to this as the ‘species topology’. Topology 6, by contrast, groups populations by geography: *cyd* with *mel-W*, and *tim* with *mel-E* ((*cyd, mel-W*), (*tim, mel-E*)). We refer to this as the ‘geography topology’. We therefore hypothesise that the history of these species can be modelled as a branching process following the species topology, but with considerable variation in lineage sorting as well as introgression that increases the rate of coalescence between sympatric populations from distinct species, as in the geography topology. Across the autosomes, the next two most highly weighted topologies (14 and 5) further support this hypothesis, as they each group one pair of sympatric taxa but otherwise match the species topology (Figure 2A).

**Figure 2.**
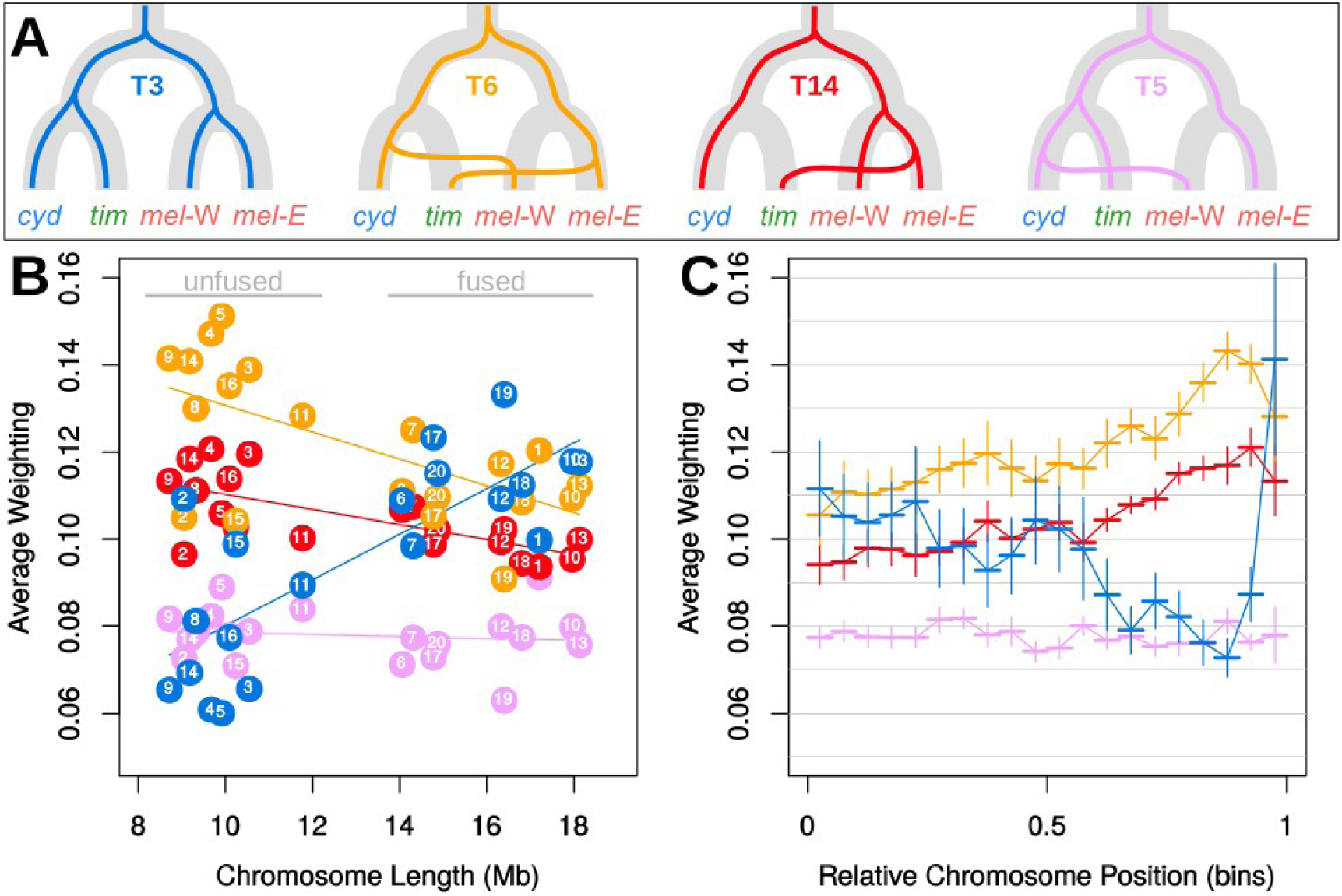
Species relationships vary consistently both among and within chromosomes. **A.** The four topologies that are most abundant across autosomes, the species topology (3, blue), the geography topology (6, yellow), and two other topologies consistent with introgression (14, red; and 5, pink). Note that topologies 14 and 5 suggest a likely predominant direction of introgression from *cyd* into *mel-W* and from *mel-E* into *tim*, and therefore the same directions are indicated in the illustration of topology 6. **B.** The average weighting for the same four topologies (colours as in **A**) for each of the 20 autosomes, plotted against the physical length of the chromosome. **C.** Average weightings for the same four topologies (colours as in **A** and **B**) binned according to their relative chromosome position, from the centre (0) to the periphery (1). Each bin represents 5% of the chromosome arm, with the range indicated by a horizontal line. Vertical lines indicate +/-one standard error.

Relative coalescence times in each genealogy provide further support for the hypothesis that introgression has led to increased clustering by geography. Topologies grouping sympatric non-sister taxa tend to have shallower splits, consistent with recent coalescence resulting from post-speciation gene flow (Figure S3). Topologies grouping allopatric non-sister taxa tend to have deeper splits, consistent with coalescence in the ancestral population and incomplete lineage sorting (Figure S3). These average branch lengths reflect a combination of both recent and ancient coalescence events and depend on population size, and should therefore be interpreted with caution. Nonetheless, the differences between topologies are in agreement with our model of recent or ongoing gene flow between both sympatric species pairs.

Our sampling design allows us to make inferences about biases in the direction of gene flow. The third most abundant topology genome-wide (Topology 14) has *tim* nested within the *mel* clade, suggesting introgression from *mel-E* into *tim* (Figure 2A). This would be expected given the much smaller range and lower effective population size of *tim*, which also has the lowest nucleotide diversity of all the taxa studied [1]. Likewise, Topology 5, which is the fourth most abundant across autosomes suggests that most introgression in the west of the Andes has been from *cyd* into *mel-W* (Figure 2A). This direction was also inferred to be the most likely in a previous study using coalescent modelling [39], and is consistent with the fact that F1 hybrids show mate preference for *H. melpomene* in experiments with Panama populations [27].

### Species relationships vary across the genome

Topology weightings vary considerably across the genome. To highlight this heterogeneity, we first focus on the two most abundant patterns of relatedness: the species and geography topologies (Figure 1 C,D). The species topology has the highest weighting in narrow peaks on some of the autosomes and across the Z chromosome, while the geography topology has a higher weighting than the species topology throughout the rest of the genome. In other words, throughout large parts of the genome, samples of *mel-W* and *mel-E* tend to be more closely related to their respective sympatric counterparts, *cyd* and *tim*, than to one another. However, the species topology tends to occur in sharp peaks, which frequently have a weighting approaching 1, indicating complete lineage sorting (Figure S4, note that these narrow peaks are not visible in Figure 1D due to smoothing). By contrast, all other topologies, including the geography topology, seldom approach a weighting of 1, indicating that they tend to be incompletely sorted, with individuals from each group being more dispersed in each genealogy. The much higher occurrence of complete sorting in the species topology is consistent with selection maintaining strong barriers between species in certain parts of the genome.

There are also strong trends in the abundance of the species and geography topologies across the 20 autosomes. All species in this clade have ten short and ten long autosomes. The latter formed through ten independent fusions in the ancestor of *Heliconius*, which had 30 autosomes [48,49].

The species topology is less abundant on the ten short autosomes as compared to the ten long, fused autosomes, and there is a fairly linear increase in its weighting with chromosome length (Figure 2A). By contrast, the geography topology shows decreasing abundance with chromosome length, and tends to be far more abundant on the short chromosomes. There is also a fairly consistent within-chromosome trend, with higher weightings for the geography topology and lower weightings for the species topology toward the outer third of the chromosomes compared to the chromosome centres (Figure 2C).

The above trends might be partly explained if the variance in lineage sorting is greatest on short chromosomes and away from chromosome centres. Indeed, we previously found a negative correlation between chromosome length and effective population size in *H. melpomene* [45]. However, several observations suggest that these patterns also reflect variation in the extent of introgression between genomic regions. Topology 14, which is consistent with introgression between *mel-E* and *tim* is less abundant than the species topology on long chromosomes and at chromosome centres, but is more abundant on short chromosomes and in chromosome peripheries (Figure 2B,C). Such a switch in rank is not expected if the short chromosomes and peripheries simply experience more variation in lineage sorting, but is consistent with differences in the extent of introgression between short and long chromosomes and between centres and peripheries. Interestingly topology 5, which is consistent with introgression between *cyd* and *mel-W*, does not show any clear relationship with chromosome length or relative chromosome position (Figure 2B,C). This implies that there may be less consistent variation in the extent of introgression between *cyd* and *mel-W* in different regions, and also further strengthens the argument that differences in the level of variation in lineage sorting alone cannot explain the differences in relatedness among chromosome regions. The chromosome-level trends for all 15 topologies are shown in Figure S5. In summary, topology weighting reveals quantitative variation in species relationships both within and among chromosomes consistent with heterogeneity in the level of introgression. However, topology weighting does not explicitly distinguish between introgression and shared ancestral variation. We therefore set out to explicitly test the hypotheses that (1) there is heterogeneity in the level of admixture across the genome, and (2) that this heterogeneity can be explained by variation in the strength of selection against introgression.

### Heterogeneous admixture suggests variable selection against introgression

We used the summary statistic *f*_d_ [15] to quantify admixture separately between *cyd* and *mel-W* and between *tim* and *mel-E*. This approach also measures an excess of genealogical clustering of sympatric non-sister taxa. However, *f*_d_ provides a normalised measure that is approximately proportional to the effective migration rate [15]. Building on previous work, we first investigated the degree to which *f*_d_ might be influenced by variation in effective population size (*N*_e_) across the genome. *N*_e_ tends to be reduced in regions of reduced recombination rate due to linked selection. By means of simulations, we find that, across a large range of realistic population sizes, *f*_d_ is a reliable estimator of admixture. Furthermore, *f*_d_ outperforms the commonly-used divergence statistics *F*_ST_ and *d*_XY_, which are both highly sensitive to *N*_e_ (Figure S6). When population sizes are very large, *f*_d_ tends to underestimate the true level of admixture. This is caused by a loss of information when population sizes are large relative to the split times: the lack of lineage sorting means that there is insufficient information available to accurately quantify admixture. The population sizes for which this is relevant are at the upper end of estimates for these species [45]. Moreover, this error would cause a conservative bias in our results, as we expect reduced admixture in low-recombination regions, where *N*_e_ is expected to be the smallest. Most important for our subsequent analysis, high background selection in regions of low recombination, which is known to influence measures such as *F*_ST_ is not likely to strongly bias our estimates using *f*_d_. We therefore conclude that *f*_d_ provides a suitable, albeit conservative, measure to test the hypothesis that species barriers are enhanced in regions of reduced recombination rate.

Computation of *f*_d_ requires the use of a ‘control’ population that is ideally allopatric and unaffected by introgression. To confirm the robustness of our results, we computed *f*_d_ with several different sets of populations, varying the control population, as well as splitting or joining each of *cyd*, *tim*, *mel-W* and *mel-E* into their two constituent sub-populations (Figure S7).

Patterns of admixture estimated by *f*_d_ show considerable heterogeneity across the genome (Figure 3A). As expected, admixture is minimal across the Z chromosome in both pairs, indicating a strong barrier to introgression. There is also heterogeneity in admixture proportion across the autosomes. This is most striking between *tim* and *mel*-E, where some regions exhibit deep troughs, implying strong, localised species barriers. Some of this heterogeneity likely reflects individual barrier loci of large effect. Indeed, the known wing-patterning loci provide a useful example. The pattern differences between *cyd* and *mel-W* are determined by regulatory modules around three major genes: *wnt-A* (chromosome 10), *cortex* (chromosome 15) and *optix* (chromosome 18) [42,50–54]. These probably act as strong barriers to introgression between *cyd* and *mel-W*, due to increased predation against hybrids with intermediate wing patterns [36]. By contrast, the shared wing patterns of *tim* and *mel*-E are thought to result from adaptive-introgression of wing patterning alleles. As expected, there is a strong reduction in admixture between *cyd* and *mel*-W in the vicinity of all three genes, while there are peaks of admixture between the co-mimetic *tim* and *mel*-*E* populations in the corresponding regions (Figure S8).

### Admixture proportions are correlated with recombination rate

We hypothesized that many loci across the genome contribute to the species barriers, which leads to the expectation that the level of admixture will be correlated with the recombination rate [16]. We quantified variation in recombination rate across the genome using high-resolution linkage maps [43] as well as using LDHelmet, which estimates the population recombination rate (ρ) based on linkage-disequilibrium (LD) in the genomic data from natural populations. At a broad scale, the map-based estimates are highly concordant with the population-based estimates, and the latter are also strongly conserved across the different species (Figure 3B, S9). There is considerable variation in recombination rate across the genome, allowing us to investigate whether admixture proportions are correlated with recombination rate.

**Figure 3.**
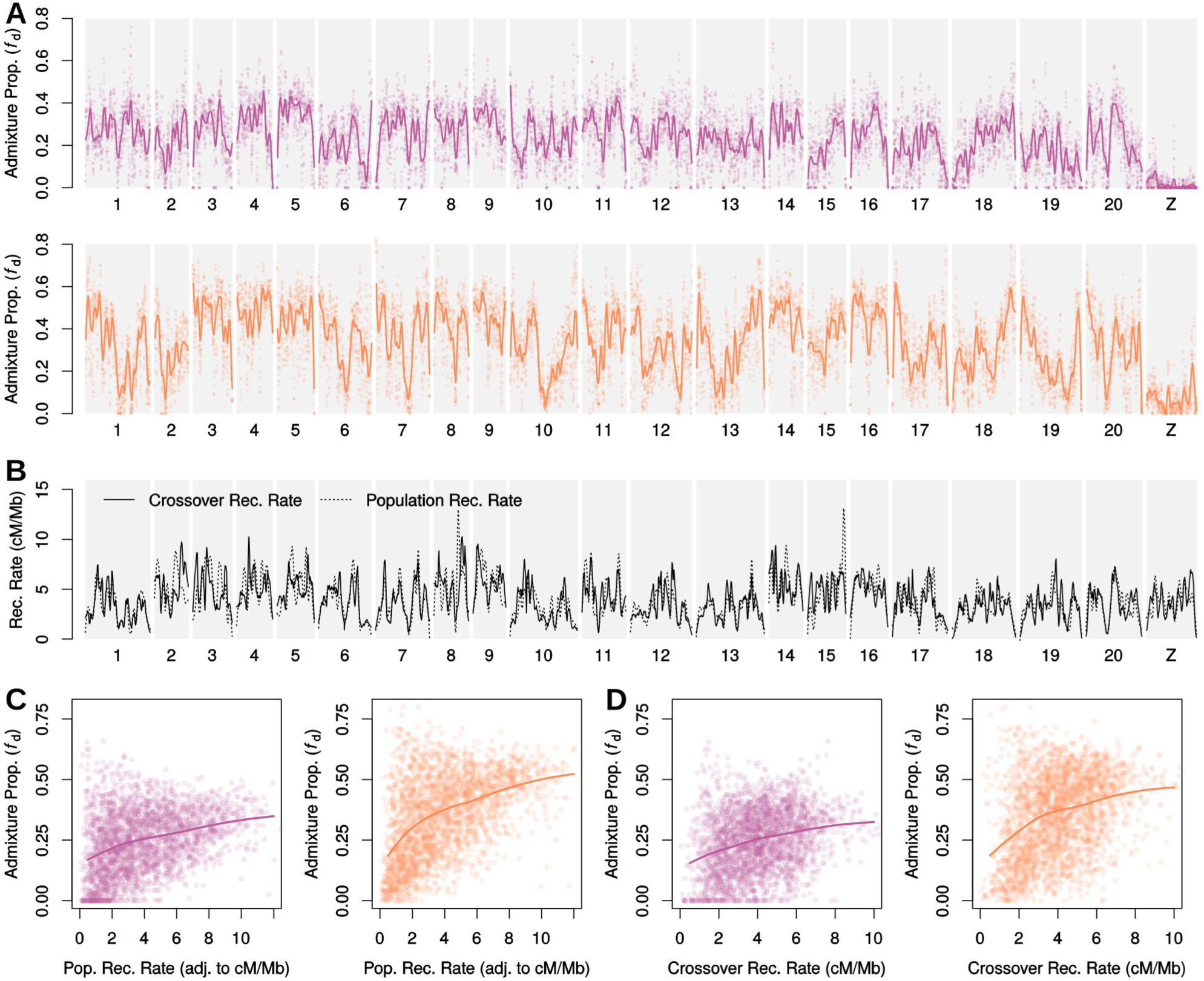
Admixture proportions are correlated with recombination rate. **A.** Estimated admixture proportions (*f*_d_) between *cyd* and *mel-W* (upper) and between *tim* and *mel-E* (lower) plotted across all 21 chromosomes in 100 kb windows, sliding in increments of 20 kb. A locally-weighted average (loess span=2 Mb) is included. Results shown are for population Sets 1 and 4 of Figure S7. **B.** Recombination rate estimated from the crossover rate in linkage maps (solid line) and the population recombination rate averaged across the four populations considered and plotted as a locally-weighted average (loess span = 2 Mb) (dashed line). See Figure S9 for a more detailed plot. **C.** Admixture proportions for *cyd* and *mel-W* (left) and *tim* and *mel-E* (right) for non-overlapping 100 kb windows plotted against the population recombination rate. Solid lines indicate the locally-weighted average (loess span = 0.75). Dashed lines indicate the same, but when windows in the outer third of each chromosome are excluded. **D.** As in C, except that the x-axis is the crossover recombination rate inferred using linkage maps.

There is a strong positive and non-linear relationship between admixture proportion and recombination rate in both species pairs (Figure 3C, 3D, S11). Strong reductions in admixture, implying barriers to introgression, are concentrated in genomic regions where recombination rates are below 2 cM/Mb. However, there is also more variability in admixture proportions in these low-recombination regions, with some showing high levels of admixture (Figure 3C, 3D). This might imply that some regions do not harbour loci that contribute to the species barrier, although the variance in admixture proportions may also be increased in low-recombination regions due to increased genetic drift resulting from enhanced linked selection. In addition, the relationship between admixture and recombination rate is clearest in the *tim* and *mel-E* pair implying that the more heterogeneous pattern of admixture across the genome between this pair is more consistent with a model in which barrier loci are widespread and recombination rate modulates the strength of the barrier to introgression.

Admixture proportions are less well predicted by the map-based estimates of crossover recombination rate (Figure 3D) compared to the inferred population recombination rate (ρ) (Figure 3C,). This probably partly reflects inaccuracy in fine-scale recombination rate estimated from the linkage maps. However, it may also be that ρ (=4*N*_*e*_*r*) provides a more meaningful predictor for the admixture proportion, as it is a composite of the per-generation recombination rate (*r*) and local effective population size (*N*_*e*_). Due to linked selection, parts of the genome with a low recombination rate and a high density of selected sites are expected to have locally reduced *N*_*e*_ and therefore reduced ρ. Indeed, ρ is strongly negatively correlated with gene density (linear regression, r^2^=0.368, p=9.67×10^−273^, Figure S10). However, there is also a weaker but significant negative relationship between gene density and the crossover recombination rate (linear regression, r^2^=0.064, p=7.93×10^−41^, Figure S10). This implies that linked selection in regions of low recombination rate may be further enhanced by a higher density of selected loci. As the conditions that enhance linked selection are the same as those expected to strengthen barriers to introgression (i.e. a high ratio of selected loci relative to the recombination rate, also called the ‘selection density’ [16]) it is to be expected that ρ would provide a better predictor of barrier strength and therefore admixture proportion. As expected, there is a negative relationship between admixture proportion and the proportion of coding sequence per window (referred to as ‘gene density’ below) (Figure S12). However, the fact that regions with a high gene density also tend to have lower recombination rates makes it difficult to determine whether such regions harbour a higher physical density of barrier loci, but this seems likely given the arguments above.

The above trends are robust to using different allopatric ‘control’ populations when estimating admixture proportions (Figure S11, S12), with the exception that using very closely related control populations lead to very low estimated rates of admixture, for which the relationships with recombination rate and gene density are not clear. See Figure S11 and Figure S12 for details.

### Large scale trends in patterns of admixture

Average chromosomal admixture proportions are negatively correlated with chromosome length (Figure 4A, 4B). This is expected given the extremely strong negative correlation between physical chromosome length (in base pairs) and average recombination rate (Figure 4C). By contrast, there is no clear relationship between chromosome length and gene density (Figure 4D). The broadly enhanced barrier to introgression on long chromosomes is therefore more consistent with an effect of increased linkage, rather than an increased density of barrier loci. As in the trends above, the relationship with chromosome length is stronger for admixture between *tim* and *mel-E* (correlation coefficient = −0.76, p=8e-05, df=18), than for admixture between *cyd* and *mel-W* (correlation coefficient = −0.52, p=0.018, df=18). Between *tim* and *mel-E*, the shortest chromosomes experience about 50% more admixture than the longest chromosomes, with the exception of chromosome 2, which has strongly reduced admixture compared to other short chromosomes with similarly high recombination rates. This might reflect a higher density of barrier loci on this chromosome, which seems possible as it also has the highest gene density of all chromosomes (Figure 4D).

**Figure 4.**
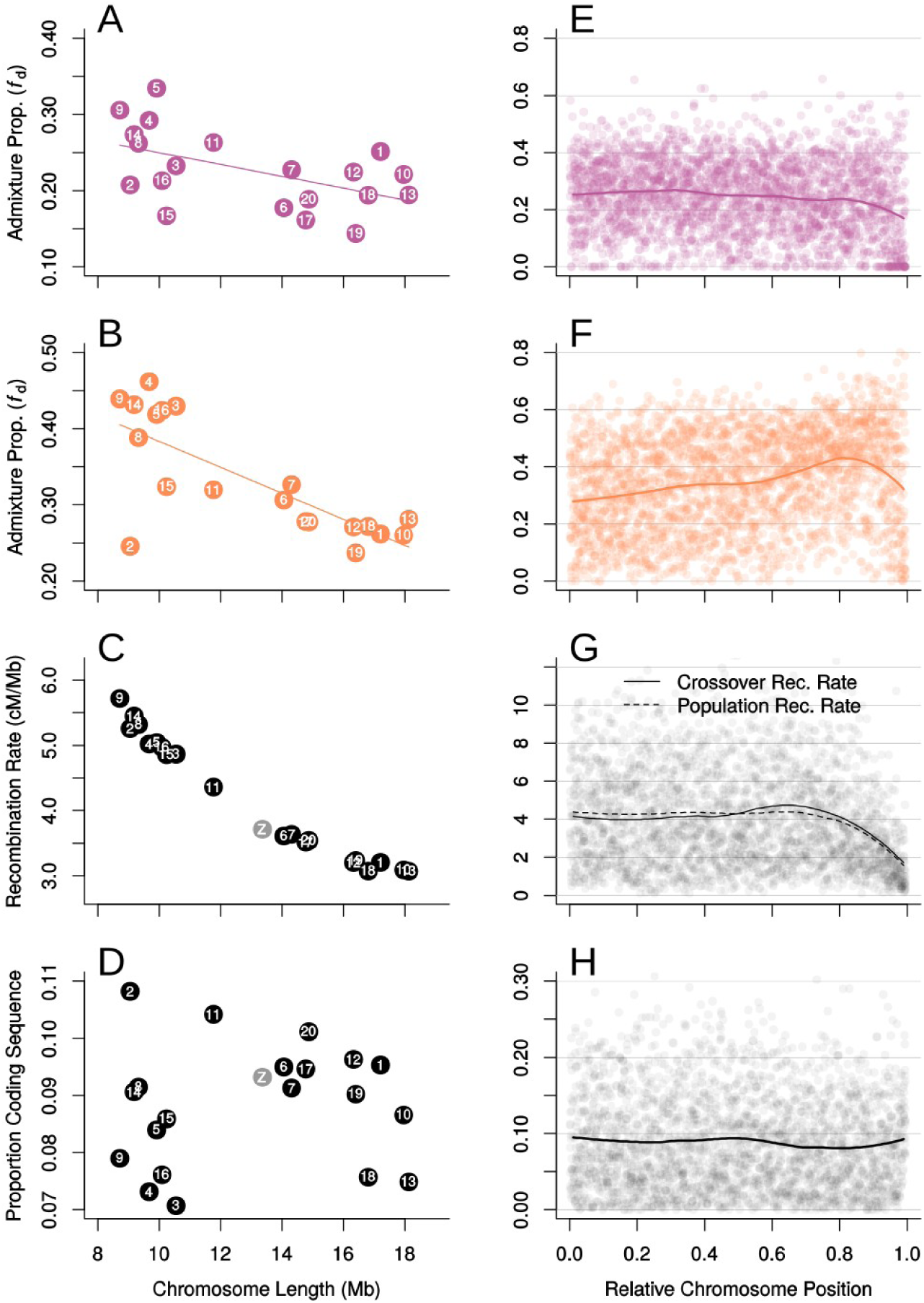
Variation in admixture proportions among and within chromosomes is explained by recombination rate and proximity to chromosome edges. **A. & B.** Estimated admixture proportions (*f*_d_) between *cyd* and *mel-W* (**A**) and between *tim* and *mel-E* (**B**) for each chromosome plotted against chromosome length. **C.** Crossover recombination rate for each chromosome plotted against chromosome length. **D**. Proportion of coding sequence for each chromosome plotted against chromosome length. **E. & F.** Estimated admixture proportions (*f*_d_) between *cyd* and *mel-W* (**E**) and between *tim* and *mel-E* (**F**) for 100 kb windows plotted against relative chromosome position. A locally-weighted average (loess span=0.25) is included. **G.** Crossover recombination rate plotted against relative chromosome position. A locally weighted average is included, along with the corresponding line for the population recombination rate (ρ). **H.** Proportion of coding sequence per 100 Kb window plotted against average chromosome position, again with a locally-weighted average shown.

As indicated by topology weighting, there is an effect of position along the chromosome on the proportion of admixture between *mel-E* and *tim*, where admixture increases on average towards the distal region of the chromosome, but decreases again at chromosome ends (Figure 4F). This is not seen in the proportion of admixture between *cyd* and *mel-W* (Figure 4E). Unlike in many other taxa, there is no consistent decrease in recombination rate toward chromosome centres. By contrast, both the crossover recombination rate and ρ show a sharp decrease at the chromosome ends (Figure 4G). Gene density is roughly uniform across chromosomes on average (Figure 4H). Theory predicts that, given a uniform recombination rate and distribution of selected loci, species barriers should weaken toward chromosome ends, leading to increased admixture [22]. Therefore, the different trends seen in the two species pairs might imply a different balance between this edge effect, which should weaken species barriers, and reduced recombination, which should strengthen them.

## DISCUSSION

Introgression effectively acts to rewrite the evolutionary history of the genome. Genome-scale data have revealed that the extent of introgression in some species may be far greater than previously imagined [1,2,55,56]. Despite the strong behavioural isolation among the three species studied here, we find that relationships among them vary dramatically across the genome, and that in some parts introgression has overwhelmed the genealogical footprints of the original population branching pattern. Similar dramatic heterogeneity in species relationships has been described in several other taxa [2,57]. For example, introgression among some *Anopheles* mosquitos has almost entirely eliminated the signal of the original species branching events on autosomes [2]. In the present study, by analysing genomes from multiple samples per population, we show that *Heliconius* species relationships vary quantitatively within and among chromosomes. Our main finding is that this variation in species relationships is predictable, and can be explained by quantitative variation in the strength of the selective barrier to introgression, which depends on the local recombination rate. Our findings therefore show how hybridisation and natural selection act in combination to shape the tree of life.

There has been considerable interest in making inferences about species barriers from the genomic landscape of divergence between hybridising species, based on the idea that selection should resist genetic homogenisation by gene flow at barrier loci. However there has also been an increasing realisation that patterns of differentiation and divergence can be influenced by unrelated factors, such as linked selection acting within species [13,58–60]. One effect of these confounding factors is that relative measures of divergence, such as *F*_ST_, can show elevated values in regions of the genome that experience stronger linked selection, even if there is no reduction in effective migration in such regions, and indeed even when there is no gene flow at all [14]. In other words, the observation of increased *F*_ST_ in regions of the genome with reduced recombination rate is not informative about the architecture of species barriers. This is particularly problematic when considering species that hybridise rarely, as the contribution of gene flow to patterns of differentiation may be small compared to that of within-species linked selection. Previous analyses of these *Heliconius* species revealed a highly heterogeneous pattern of *F*_ST_, in which even known wing patterning loci that have a major impact on hybrid fitness are not particularly prominent [1]. Until now, it has not been known whether barrier loci are indeed widespread throughout the genome in these species.

Our results suggest that there are highly polygenic barriers that maintain these *Heliconius* species. Genome windows in regions of high recombination rate (>5 cM/Mb) almost invariably show increased levels of admixture, whereas windows showing reduced admixture are concentrated in parts of the genome with low recombination rates (<2 cM/Mb). This is consistent with theoretical expectations that barrier loci will cause a stronger localised reduction in introgression in regions of lower recombination rate [16,21–23]. Interestingly, some windows in regions of low recombination nevertheless show high levels of admixture. This may indicate that barrier loci, although abundant, are not ubiquitously distributed across the genome. However, we also expect increased variance in levels of admixture in these low-recombination regions due to increased genetic drift, which will be compounded by the reduced independence among sites in the 100 kb windows. This increased variance does not explain the positive relationship between admixture and recombination, however. Selection against introgression also produces a global pattern of decreasing admixture with chromosome length. Long chromosomes, which have similar gene density but lower per-base recombination rates than short chromosomes, form stronger barriers to introgression on average. It is likely that species barriers are also stronger in gene-rich regions, due to an increased density of barrier loci. While we did find a weak trend of reduced admixture in gene-rich regions, this is difficult to interpret, as the recombination rate is also lower in gene-rich regions. A final factor that could influence our conclusions is the fact that barrier loci may not be expected to accumulate randomly across the genome. Some models predict that, under a scenario of ecological divergence in the face of gene flow, the accumulation of barrier loci may be clustered [23,61]. This could increase the correlation between admixture and recombination, and perhaps lead to overestimation of the density of barrier loci. Clustering could theoretically be further enhanced by genomic rearrangements between species that physically suppress recombination in hybrids. However, we have previously found that there are no major inversions maintaining barriers between *H. melpomene* and *H. cydno* [43]. Nonetheless, the lack of information about the distribution of barrier loci and their effect sizes means that it is currently not possible to estimate the number of loci involved, except that it is probably very large.

In agreement with previous findings, we find that barriers to introgression are far stronger across the Z chromosome compared to autosomes. Enhanced barriers to introgression on sex chromosomes have been observed in genomic studies of a range of taxa with both XY and ZW systems [1–3,62]. This has been attributed to a more rapid build-up of incompatibilities due to hemizygozity and a key role played by sex chromosomes in reproduction and fertility. Comparing genetic differentiation on sex chromosomes to autosomes can be complicated by their reduced effective population size, and this can be further confounded by changes in population size, which can affect sex chromosomes differently [63]. However, our simulations show that these factors cannot explain the reduction in admixture detected here using *f*_d_. In these *Heliconius* species, hybrid female sterility is associated with one or more loci on the Z chromosome [35,64,65]. The observed reduction of admixture across the Z chromosome must result from selection against foreign Z chromosome alleles in backcross progeny and their descendants, such that there are opportunities for independent assortment of chromosomes prior to selection. Segregation of sterility in backcrosses has indeed been observed in crossing experiments [35,65,66]. This also means that there are opportunities for recombination before selection. The fairly even reduction in admixture we observe across most of the chromosome is therefore perhaps surprising, and implies that there are multiple barrier loci spread across the Z chromosome. A similar pattern of widespread incompatibilities throughout much of the sex chromosome has been shown experimentally between *Drosophila* species [67].

The model proposed here of a highly polygenic species barrier between *H. melpomene* and its relatives contrasts with previous studies that identified a few major effect loci that control differences in wing pattern and mate preference between *H. cydno* and *H. melpomene* [36,66]. However, multiple additional behavioural and ecological differences are known to distinguish these species [29,68], and each of these may have a more polygenic basis. Interestingly, unlike *H. cydno*, *H. timareta* races have wing patterns that commonly match those of the local *H. melpomene* races. Hence, the large effect wing patterning loci do not contribute to the barrier between *H. timareta* and *H. melpomene*. This difference might explain why admixture between this pair is more strongly correlated with recombination rate. When barrier loci are weak and dispersed across the genome, admixture proportions should be more strongly predicted by recombination rate than when there are few large-effect barrier loci. Hence, perhaps counter-intuitively, the more heterogeneous pattern of admixture between *H. timareta* and *H. melpomene* is in fact more consistent with the model of small-effect barrier loci evenly distributed across the genome. The heterogeneity reflects the underlying heterogenous recombination landscape, rather than a patchy distribution of large-effect barrier loci.

The cause of the more even pattern of admixture between *H. cydno* and western *H. melpomene* is less clear, but it may be explained by epistatic selection on the patterning loci. Lindtke and Buerkle [69] distinguish between two types of epistatic barrier loci: “DMI-type” incompatibilities that cause reduced fitness in hybrids but can be broken down by recombination in backcross progeny, following the formulation by Dobzhansky [4] and Muller [7], and “pathway-type” incompatibilities that reduce fitness in recombinant hybrids in which co-adapted alleles become separated. Simulations show that pathway-type incompatibilities can produce pronounced localised barriers to introgression, while DMI-type incompatibilities can have more even, genome-wide effects, if selection is strong enough [69]. Hybrids between *H. cydno* and *H. melpomene* have intermediate wing patterns that do not resemble the recognisable warning patterns of either species, making them roughly twice as vulnerable to predation [36]. The patterning loci may act as DMI-like incompatibilities, as backcrossing restores a recognisable warning pattern to a proportion of the progeny. It is therefore possible that the presence of a few large-effect incompatibilities could in fact explain the less heterogeneous landscape of introgression between *H. cydno* and *H. melpomene*, although this requires further investigation.

In conclusion, our findings imply that barrier loci have accumulated rapidly in the 1-1.5 million years over which these butterfly species have diverged. This joins a growing number of examples showing that selection against introgression between fairly young species can be pervasive across the genome [16,24,25]. Further work is still required to determine the generality of these trends, and also to account for complications such as clustered barrier loci and epistasis. These new insights into the polygenic nature of species boundaries highlight the dangers of assuming strictly neutral evolution when modelling speciation. Models that incorporate variable selection pressures among sites [55,70] are likely to be more realistic. Our results here are intriguing in that they show that, despite the widely distributed barriers across the genome, introgression has nonetheless dramatically reshaped species relationships. A few recent examples have shown how introgression can lead to dramatically different topologies across genome regions, but our data goes further in showing (1) how this phylogenetic heterogeneity can be predicted by recombination rate and (2) how relationships can vary across a species range. The focal species *H. melpomene* has very different relationships to its sister taxa both depending on the genome region and on the population sampled. These patterns raise questions about how we view the species as an entity and the degree to which animal life can be accurately viewed as a bifurcating tree.

## METHODS AND MATERIALS

### Samples and genotyping

We used whole genome resequencing data from 92 wild-caught butterflies (Table S1) [1,42,45,52,63]. Reads were mapped to the *H. melpomene* genome assembly v2 [49] using BWA mem v0.7.2, using default parameters. Read depth was computed using GATK v3.4 DepthOfCoverage [71]. Average read depth across all 92 samples was 29.22 (Table S1). Genotyping was performed using the GATK v3.4 HaplotypeCaller and GenotypeGVCFs tools [71], using default parameters except that heterozygosity was set to 0.02. Each geographic population (10 samples each) was genotyped separately. Variant sites were accepted only if the quality (QUAL) value was ≥ 30, and individual genotype calls were accepted only where the sample depth of coverage for the position was ≥ 8. Scaffold positions in the Hmel2 assembly were converted to chromosome positions based on the most recent scaffolding [43], now released as Hmel2.5. Two sets of filtered SNPs were generated for the analyses below. In addition to the requirement of ≥ 8X depth of coverage, SNPs were required to be biallelic and sites at which more than 75% of samples were heterozygous, or where the minor allele was present in only a single haploid copy, were discarded. SNP Set 1 had the further requirement that at least nine out of the ten samples representing each of the nine ingroup populations, and one of the two outgroup samples, had an accepted genotype call, resulting in 14,406,386 SNPs. SNP Set 2 had the less stringent requirement that at least four of the samples from each ingroup population, and one of the outgroup samples, had an accepted genotype call, resulting in 23,084,596 SNPs.

### Principal Components Analysis

We used Eigenstrat SmartPCA [72] to investigate population structure and confirm sample identity. SNP Set 1 was used for this analysis.

### Topology Weighting

To quantify genealogical relationships among taxa, we used topology weighting by iterative sampling of subtrees (*Twisst*) [46] (github.com/simonhmartin/twisst). This also made use of SNP Set 1. Genotypes for all samples were first phased and imputed using SHAPEIT v2 [73,74]. Neighbour-joining phylogenies were inferred for windows of 50 SNPs, following extensive simulations by Martin et al. [46]. Exact weightings were computed for all inferred genealogies that could be simplified to ≤ 2000 remaining sample combinations (see reference [46] for details). In cases where this was not possible, approximate weightings were computed by randomly sampling combinations of haplotypes until estimated weightings for all 15 possible topologies had a 95% binomial confidence interval of <0.05. Confidence intervals were computed according to the Wilson method, as implemented by the package binom [75] in R [76].

### Admixture proportions

We estimated admixture proportions for 100 kb windows using *f*_d_ [15], which is based on the so-called ABBA-BABA test [77,78]. This analysis was implemented using the python script fourPopWindows.py, available from github.com/simonhmartin/genomics_general. To ensure that *f*_d_ is not affected by confounding factors such as effective population size and selective sweeps, we first tested its performance in quantifying the proportion of admixture using coalescent simulations, and compared it to other methods used to study admixture and genomic divergence. We used msms [79] to simulate the evolution of independent windows of 50 kb, with a population recombination at rate of 1%. The models used, along with the range of effective population sizes and rates of gene flow tested are shown in Figure S6.

Analyses of real data focused on quantifying admixture between the two sympatric species pairs: *cyd* and *mel-W*, and *tim* and *mel-E*. *f*_d_ was computed using a range of combinations, including different allopatric control populations and either combining the two races that represent each broad geographic area (‘East’, ‘West’ and ‘Guiana’) or keeping them separate (Figure S7). SNP Set 2 was used for these analyses, with the added requirement that for the given run, at least 50% of samples in each population were genotyped and the outgroup was fixed for the ancestral state.

### Recombination rate estimation

Recombination rates were estimated in two different ways. First, we used the high-resolution linkage maps recently produced for *H. melpomene*, *H. cydno* and hybrids [43] to estimate the local crossover rate. The recombination rate was computed as the slope of the locally weighted regression (loess span = 2 Mb) between physical position and map position along each chromosome [45,80]. Note that because recombination is male-limited in Lepidoptera, the values presented here represent the male-specific recombination rate. Conversion to an effective recombination rate at the population level would require knowledge of the effective sex ratio, which we don not have, so we chose here to use the male-specific rate. Second, we computed the population recombination rate (ρ) for 100 kb windows using the maximum likelihood method implemented in LDHelmet [81]. This analysis was run separately for each population of 20 samples (i.e. combining races from the same area following the results of the PCA in Figure 1), using SNP Set 1, phased as described above. A window size of 50 SNPs was used, along with the recommended range of pre-computed pairwise likelihoods. For convenience, ρ values were converted to cM/Mb by scaling values for each chromosome according to the map length of each chromosome, averaged across the three linkage maps used. As above, these values are therefore scaled to the male-specific recombination rate.

## ACKNOWLEDGEMENTS

This work was funded by ERC grant SpeciationGenetics (339873) to CDJ. SHM was funded by a research fellowship from St John’s College, Cambridge. CS was funded by Fondos Concursables Universidad del Rosario 2016-PIN-2017-001. This work made use of the Darwin Supercomputer of the University of Cambridge High Performance Computing Service (http://www.hpc.cam.ac.uk/), provided by Dell Inc. using Strategic Research Infrastructure Funding from the Higher Education Funding Council for England and funding from the Science and Technology Facilities Council. We thank Dorothea Lindtke for contributing to extensive discussions and exploration of our results, and Sarah Barker for technical assistance. We are also grateful to Richard Merrill, Claire Mérot, Markus Möst, Steven Van Belleghem and Konrad Lohse for useful discussions that shaped this paper.

## SUPPLEMENTARY INFORMATION

**Table S1.**
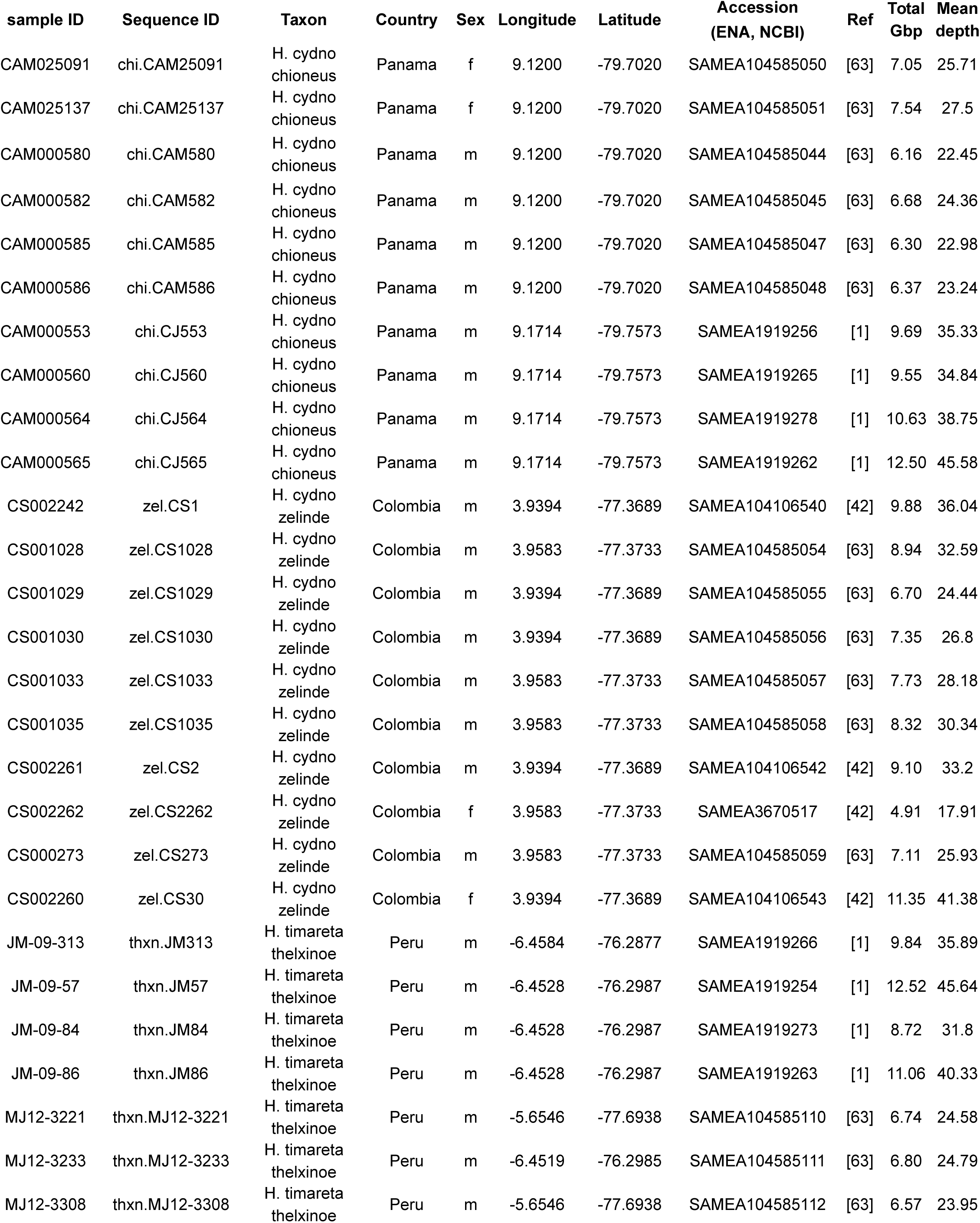
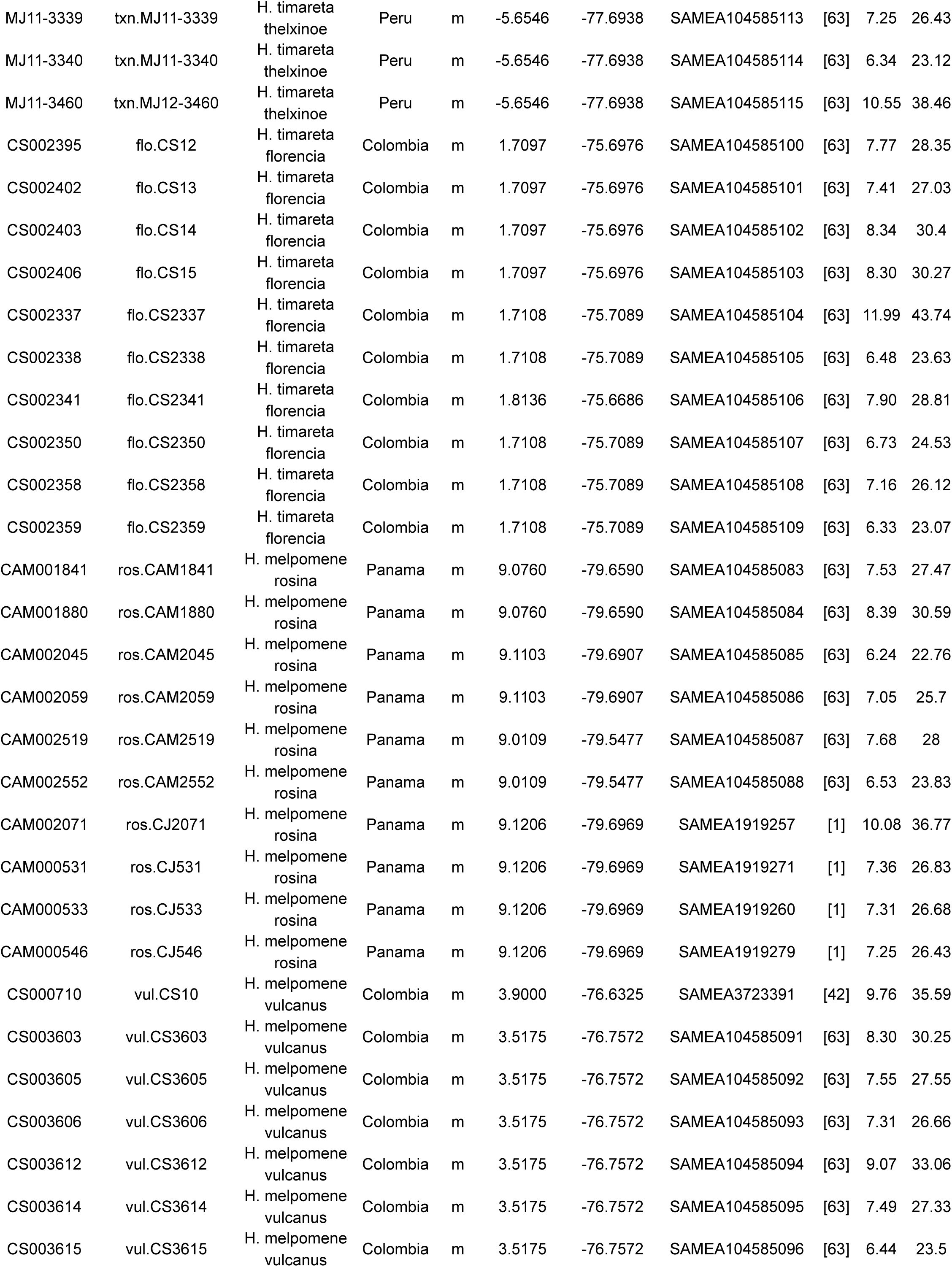
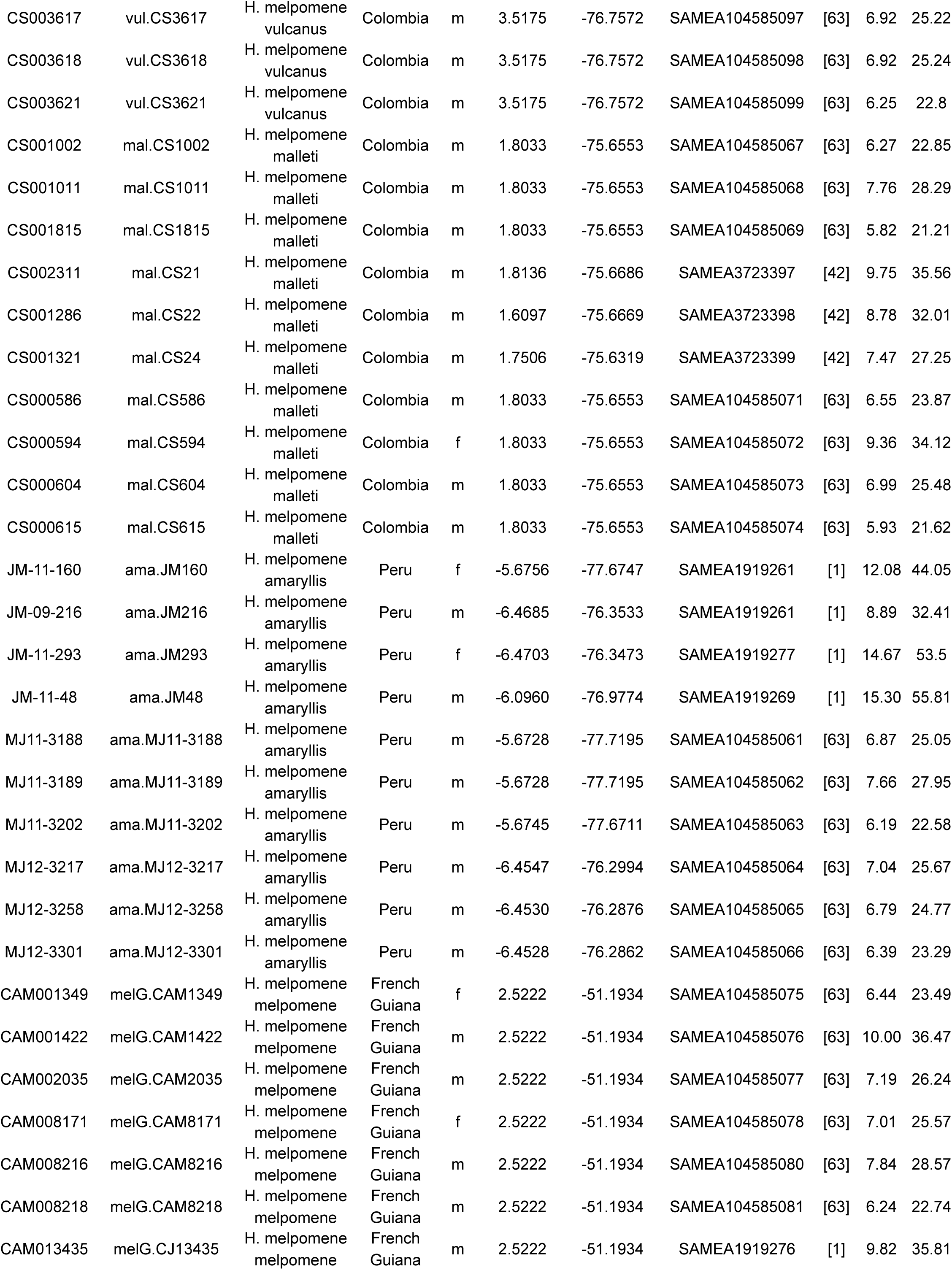
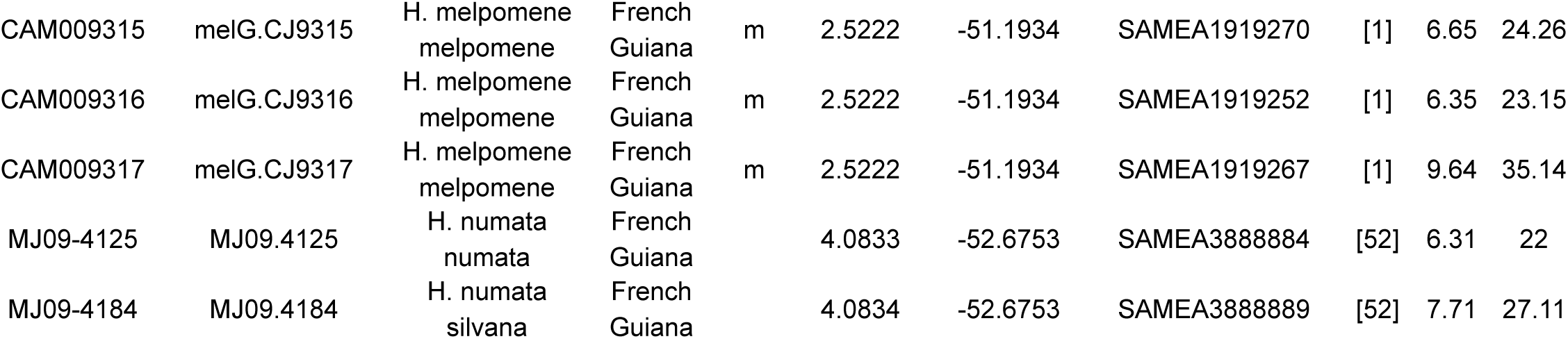
Sample and sequencing information

**Figure S1.**
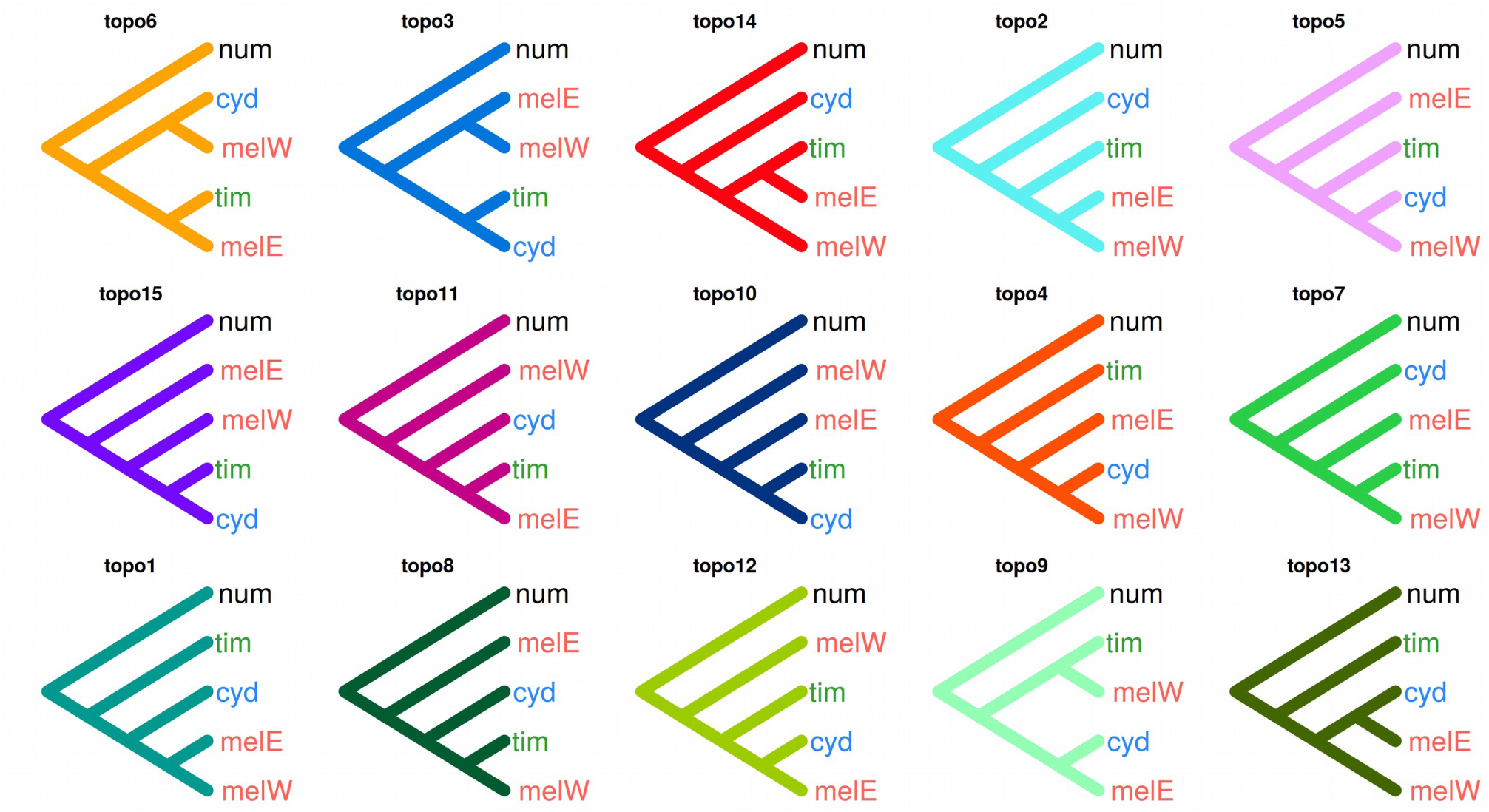
The fifteen possible topologies describing genealogical realtionships among *H. cydno* (*cyd*), *H. timareta* (*tim*), and *H melpomene* from the west (*mel-W*) and east (*mel-E*) of the Andes. The outgroup, *H. numata* (*num*), was used to polarize the genealogies. Topologies are arranged in order of their average weighting across the whole genome, from highest to lowest.

**Figure S2.**
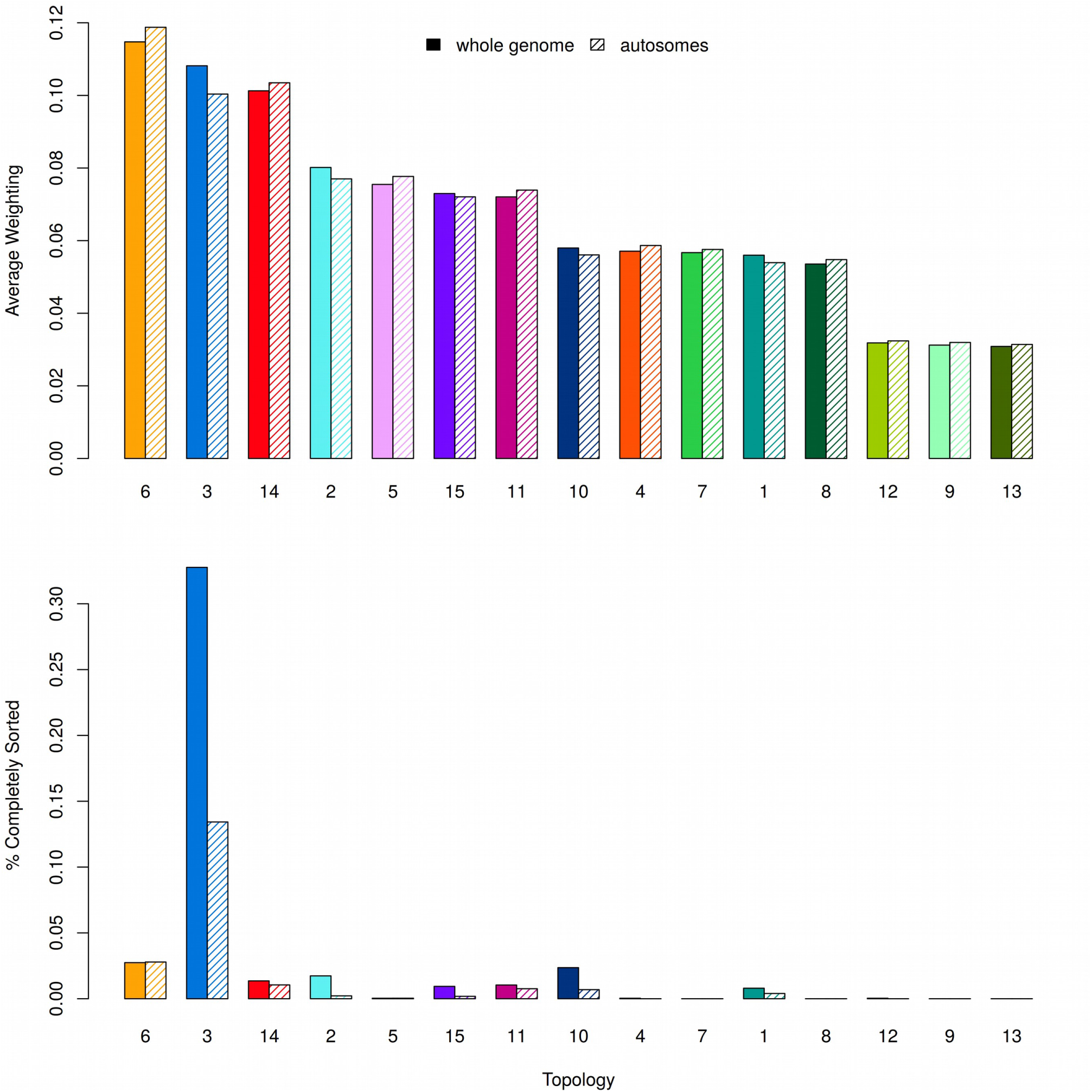
Average weightings and levels of sorting Upper: Average genome-wide weightings for the 15 topologies in Figure S1, ordered accordingly. Corresponding weightings for autosomes only are also indicated. **Lower:** The percentage of windows in which the genealogy is completely sorted (i.e. has a weighting of 1).

**Figure S3.**
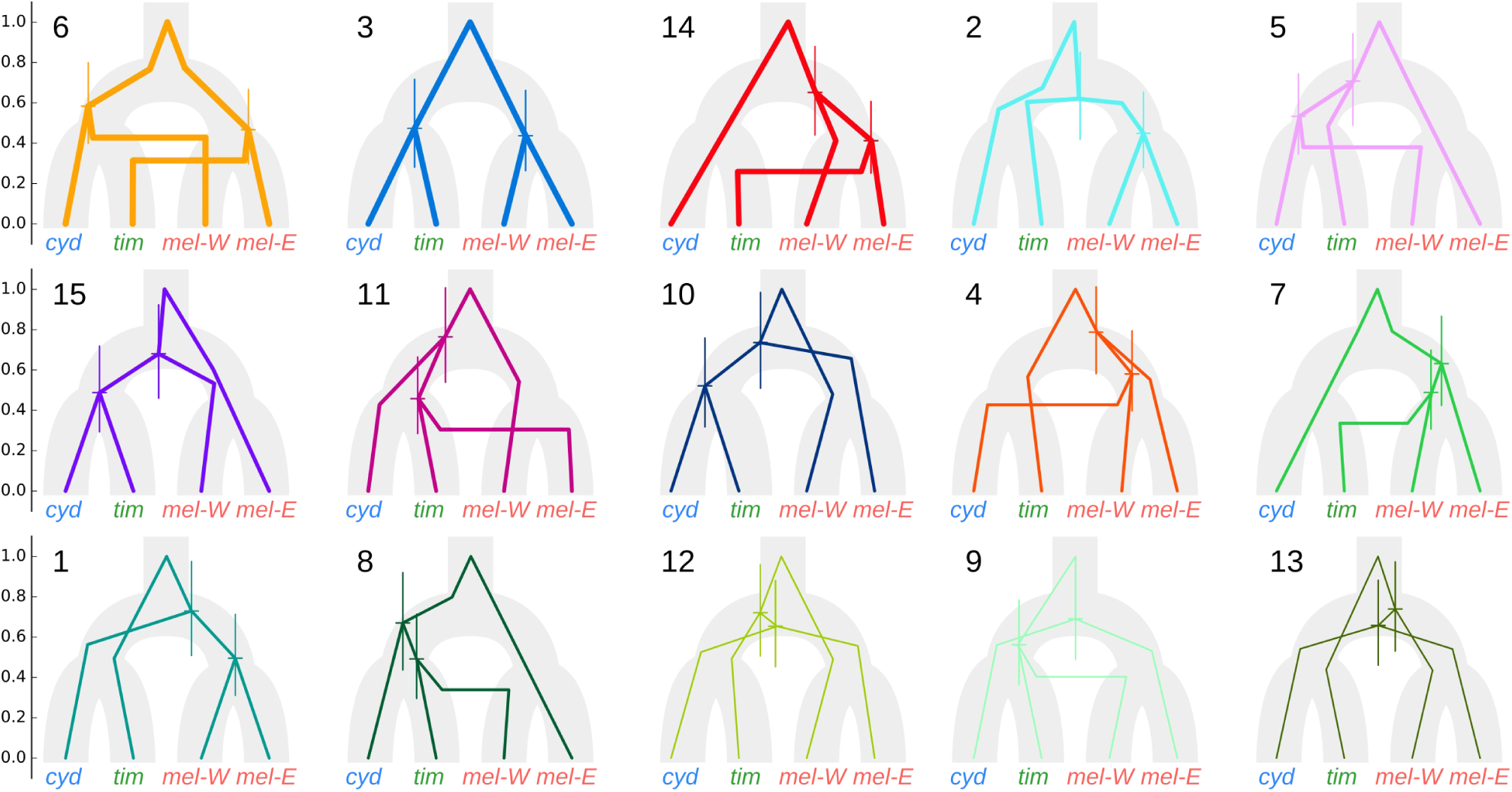
Average relative coalescence times for each topology. The fifteen possible topologies as in Figure S1, here illustrated as lineages within the presumed population branching tree. Topologies are ordered according to their average genome-wide weighting, and line widths are drawn proportionally. Coalescence times were calculated as the median branch length separating daughter taxa in each genealogy, scaled for each window by the branch length to the outgroup, to control for substitution rate variation among windows. The 25% and 75% quantiles for each split are indicated by vertical lines. Gene flow causing introgression is indicated where lineages cross between populations. Note that the each scenario shown here represents a hypothesis, and is one of several possible scenarios that can give rise to the same genealogy. These hypotheses were guided by the split times: more ancient split times are consistent with lineage sorting effects in the ancestral population (e.g. topology 13), whereas more recent split times are consistent with introgression (e.g. topology 14). It is important to note that the split times shown here represent the average across the whole genome and across multiple samples, and therefore represent the average over a range of different histories. Finally, because coalescence time will always pre-date the time of introgression, an arbitrary lag time is added prior to each introgression event. In reality, the length of this period depends on population size, and can therefore not be estimated with this technique. For this reason, these relative times of introgression between different taxa must be interpreted with caution.

**Figure S4.**
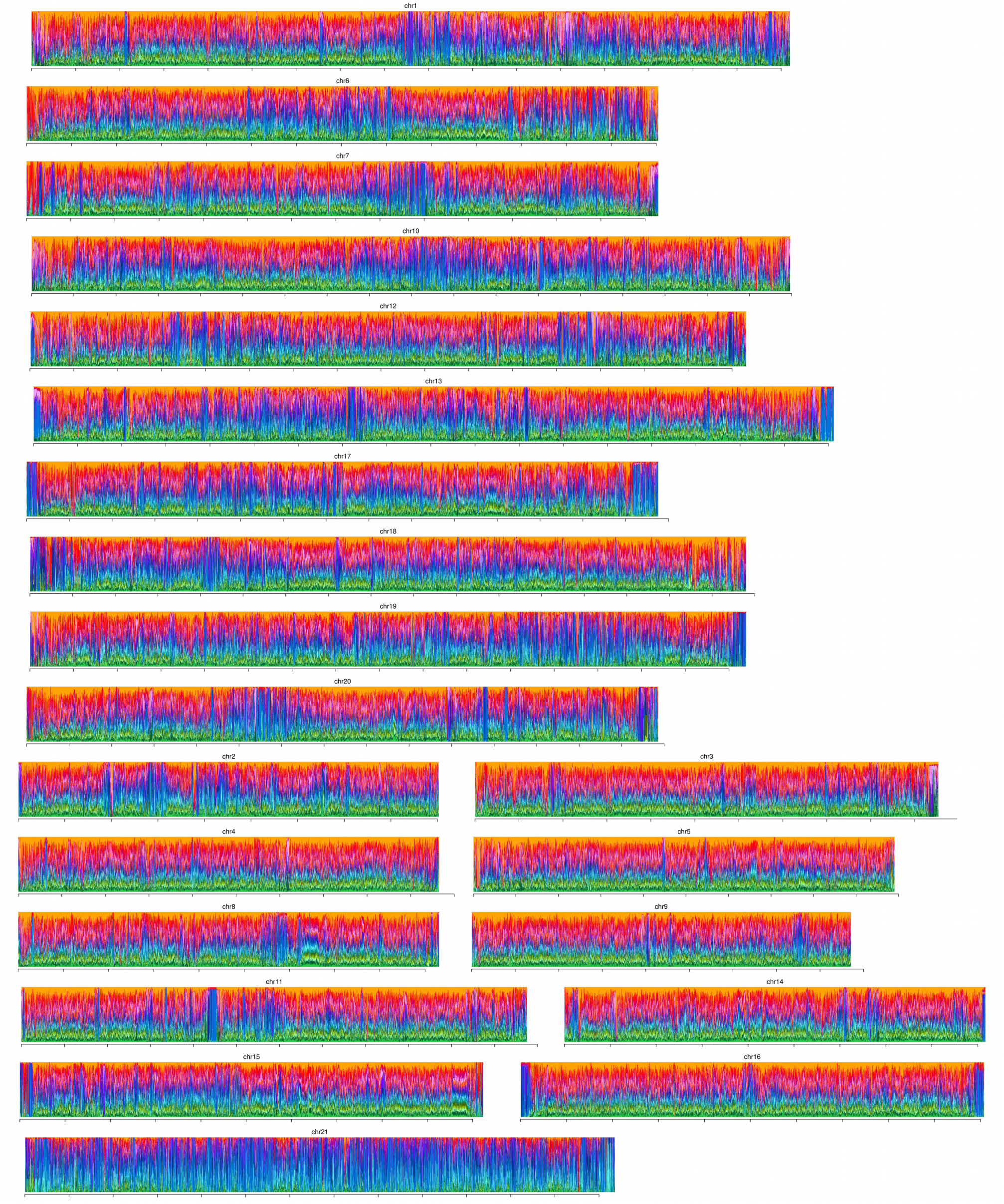
Raw weightings for all topologies across all chromosomes. See Figure S1 for the colour legend. Raw values without smoothing are plotted here, unlike in Figure 1 of the main paper. Weightings are stacked so that all 15 topologies can be distinguished. X-axis tick marks are spaced by 1 Mb.

**Figure S5.**
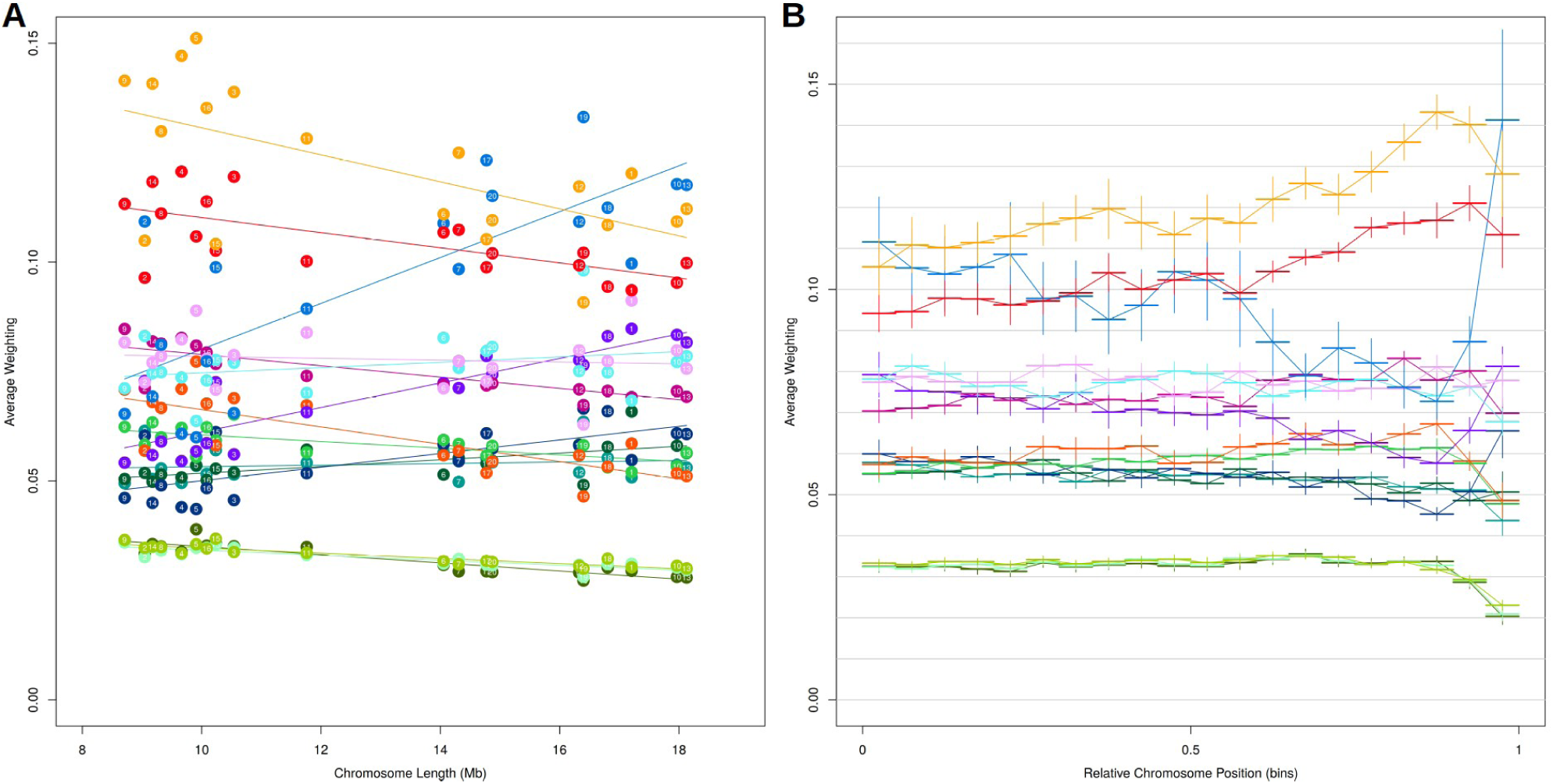
Heterogeneity in topology weightings among and within chromosomes. **A.** The average weighting for all 15 topologies (colours as in Figure S1) for each of the 20 autosomes, plotted against the physical length of the chromosome. Fitted linear regressions are shown for reference. **B.** Average weightings for all 15 topologies (coloured as in Figure S1) binned according to their relative chromosome position, from the centre (0) to the periphery (1). Each bin represents 5% of the chromosome arm, with the range indicated by a horizontal line. Vertical lines indicate +/- one standard error.

**Figure S6.**
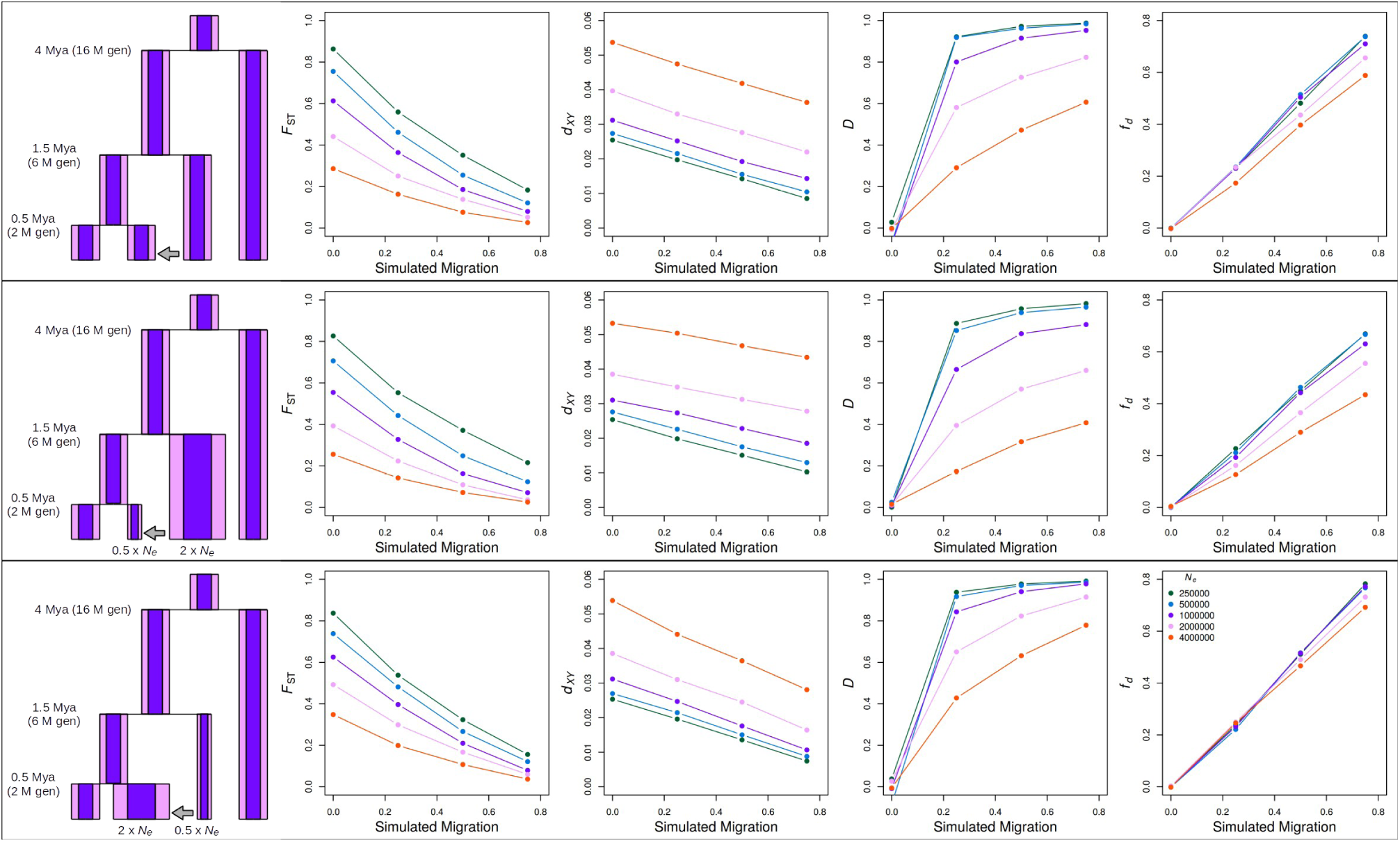
Testing the robustness of *f*_d_ to estimate the admixture proportion. Simulations show that *f*_d_ is largely robust to effective population size (*N*_e_). Sequences were simulated following the model on the left, with a range of different population sizes, indicated by different colours (two are shown in the model for example). Simulated divergence times were chosen to approximate the splits between the outgroup silvaniform clade and the clade of *H. melpomene*, *H. cydno* and *H. timareta* (~4 million years ago, [47]), and the divergence between *H. melpomene* and the ancestor of *H. cydno* and *H. timareta* (1-1.5 Mya, [38,39]). Population sizes ranging from 250,000 to 4,000,000 were tested. For comparison, other divergence and admixture statistics are included. Relative and absolute divergence statistics *F*_ST_ and *d*_XY_ are both strongly dependent on *N*_*e*_. Patterson’s D statistic is strongly affected by *N*_e_ and is non-linear. By contrast, *f*_d_ is approximately proportional to the simulated level of migration, and is largely unaffected by *N*_*e*_, except when *N*_e_ is large in which case *f*_d_ tends to underestimate the simulated admixture proportion. This is consistent with a loss of power with reduced lineage sorting in large populations. *N*_e_ for *H. melpomene* was estimated to be 2-3 Million [45], suggesting that admixture would indeed be weakly underestimated. However, as we are primarily interested in testing for reduced admixture in parts of the genome with reduced recombination rate, which usually corresponds to reduced *N*_e_ due to enhanced linked selection, the observed bias would have a conservative influence on our main analysis. We also tested simulated histories in which the donor and recipient populations undergo an expansion and contraction, respectively (second row), or the inverse (third row). Expansion of the donor population causes an exaggeration of the underestimate of admixture when *N*_e_ is large, but otherwise these changes don’t have a significant effect on the performance of *f*_*d*_.

**Figure S7.**
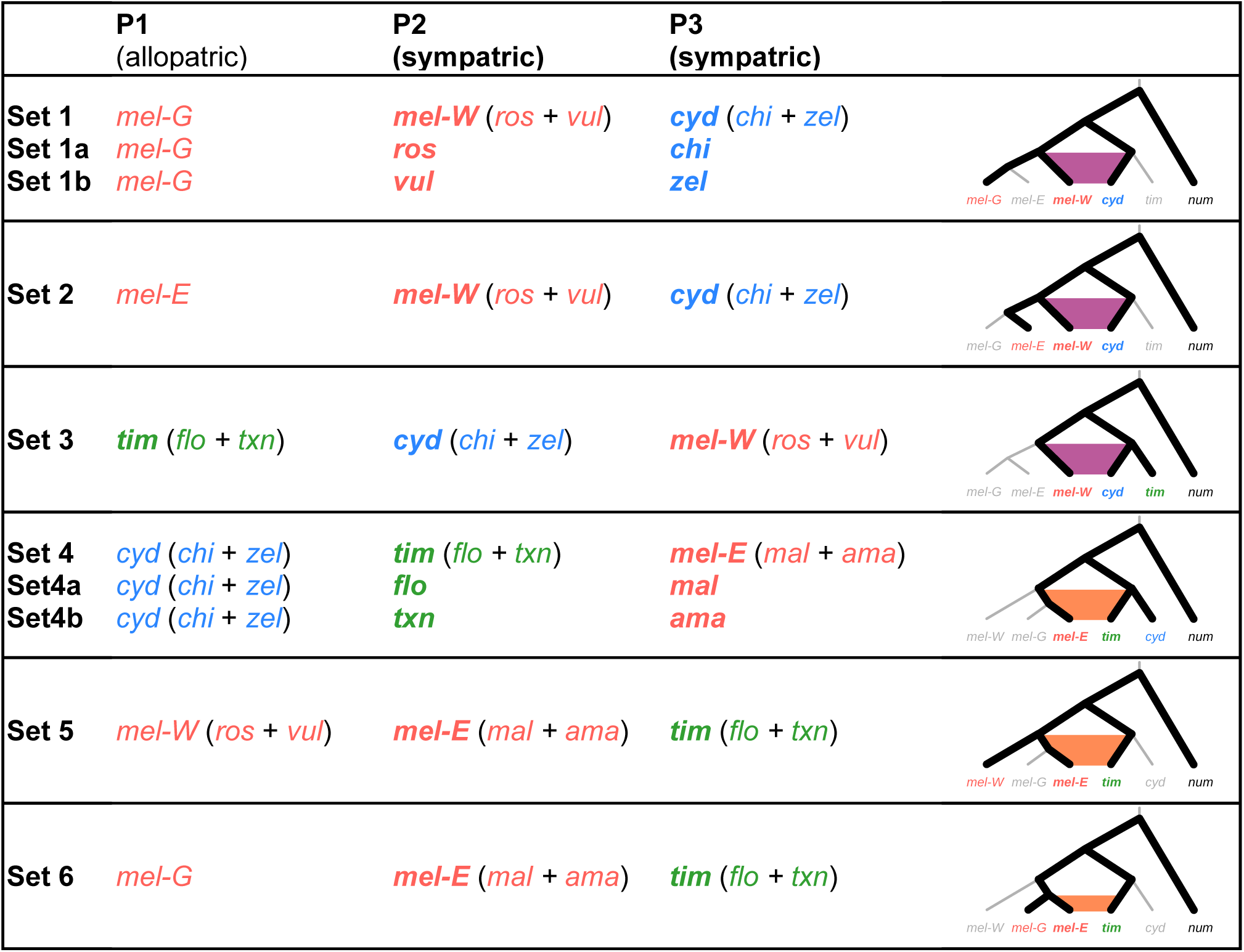
Sets of taxa used to estimate admixture proportions using *f*_d_. Sets 1-3 were used to estimate admixture between *cyd* and *mel-W*. Sets 4-6 were used to estimate admixture between *tim* and *mel-E*. In each set, P2 and P3 represent the two sympatric populations between which the level of admixture is to be measured. P1 represents the allopatric ‘control’ population that is closely related to P2, but thought not to be subject to contemporary hybridisation with P3. The figure on the right shows, for each set, the relationships among the three populations considered (bold lines), as well as the period over which admixture between P2 and P3 can be detected given P1 (shaded). In all sets, H. numata (*num*) was used as the outgroup. ‘*ros’* = *H. m. rosina*, ‘*vul’* = *H. m. vulcanus*, ‘*mal’* = *H. m. malleti*, ‘*ama’* = *H. m. amaryllis*, ‘*mel-G’* = *H. m. melpomene* from French Guiana, ‘*chi’* = *H. c. chioneus*, ‘*zel’* = *H. c. zelinde*, ‘*flo’* = *H. t. florencia*, ‘*txn’* = *H. t. thelxinoe*.

**Figure S8.**
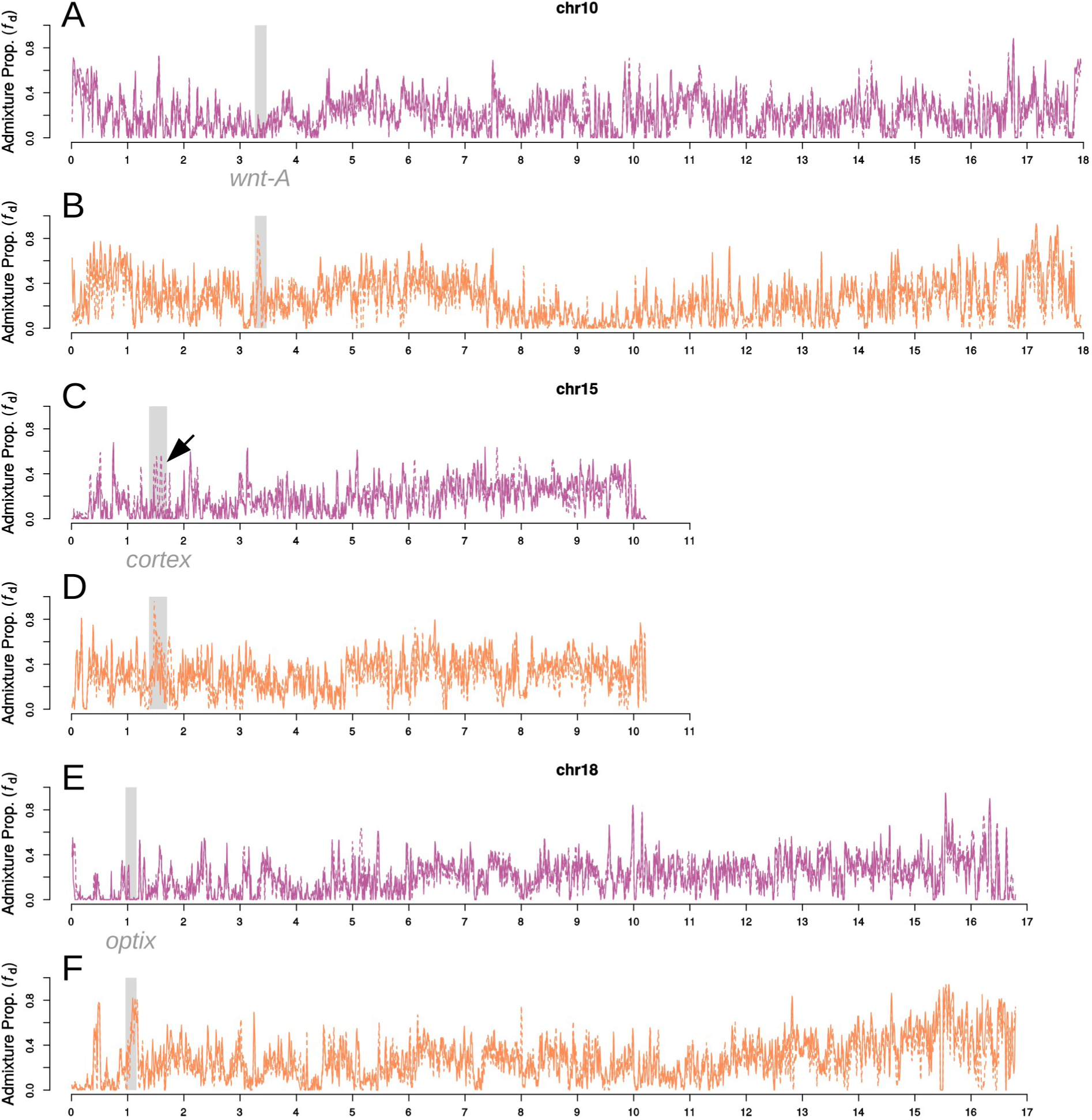
Fine scale patterns of admixture across four chromosomes containing wing patterning loci. Estimated admixture proportion (*f*_d_) computed in 20 Kb slinding windows, sliding in increments of 5 Kb, plotted across three chromosomes carrying known wing patterning genes: *wnt-A* (chromosome 10), *cortex* (chromosome 15) and *optix* (chromosome 18). For A, C, and E, *f*_d_ was computed between cyd and mel-W using Set 1a (solid purple line) and Set 1b (dashed purple) (see Figure S7). For B, D, and F, *f*_d_ was computed between *tim* and *mel-E* using Set 4a (solid purple line) and Set 4b (dashed purple). In all cases, there is reduced admixture between *cyd* and *mel-W*, and elevated admixture between *tim* and *mel-E* in the vicinity of the patterning loci, consistent with a barrier to introgression in the former, but not the latter. The one exception is the cortex locus on chromosome 15, at which there is elevated admixture for Set 1b (i.e. between *H. cydno zelinde* and *H. melpomene vulcanus*, indicated by an arrow). This has in fact been previously recorded as a probable rare instance of introgression of a wing patterning allele between *H. cydno* and *H. melpomene* [42]. This allele appears to be responsible for the dorsal melanisation of the hindwing yellow bar in *H. m. vulcanus*. Therefore, these loci provide robust support for the use of *f*_d_ to quantify admixture between these taxa.

**Figure S9.**
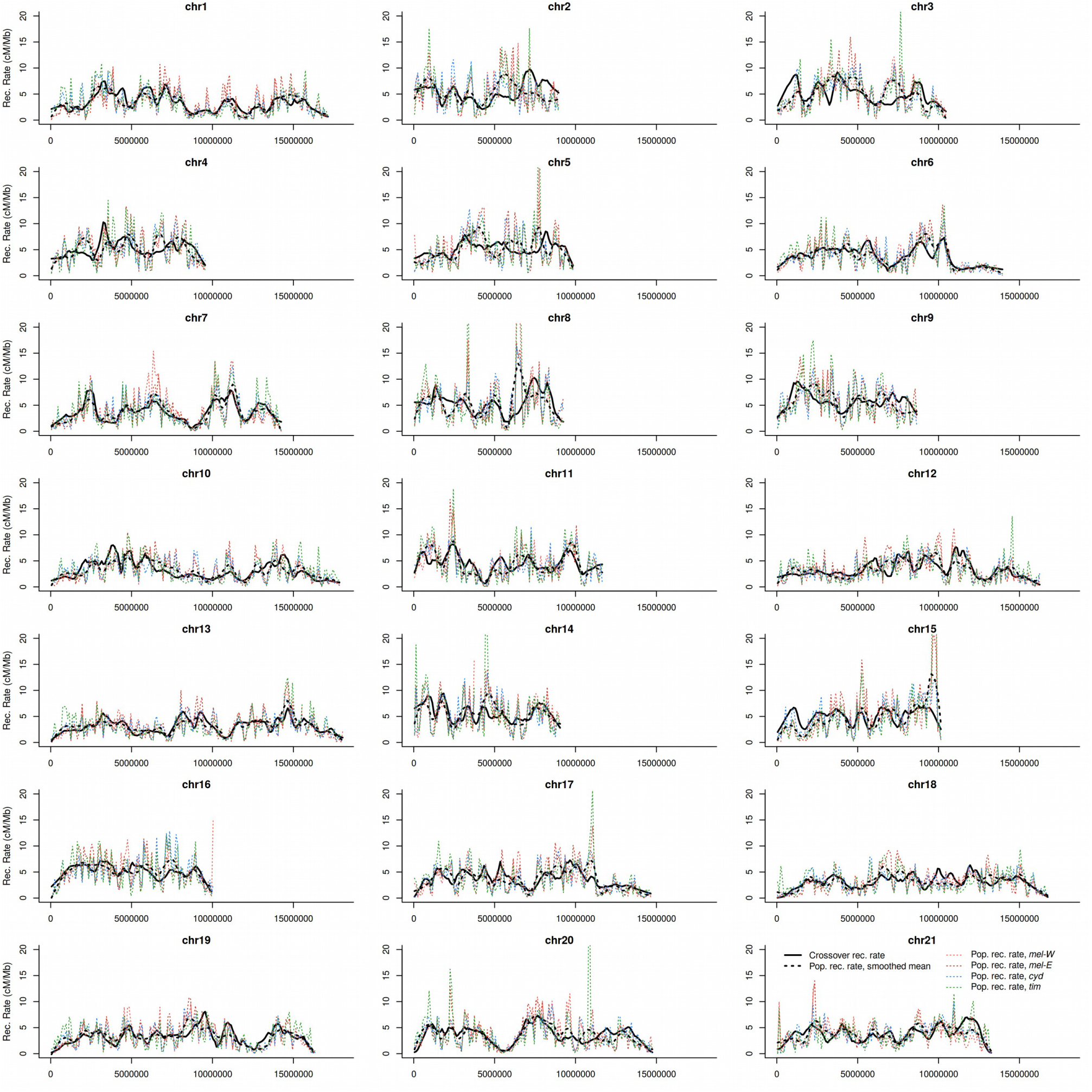
Recombination rates plotted across chromosomes. Solid lines show the crossover recombination rate estimated from linkage maps [43]. Dashed lines show the maximum likelihood estimate for the population recombination rate (ρ), computed for 100 Kb windows separately for *cyd*, *tim*, *mel-W* and *mel-E* (indicated by colours). The black dashed line indicates the mean ρ across the four populations, plotted as a locally-weighted average (loess span = 2 Mb).

**Figure S10.**
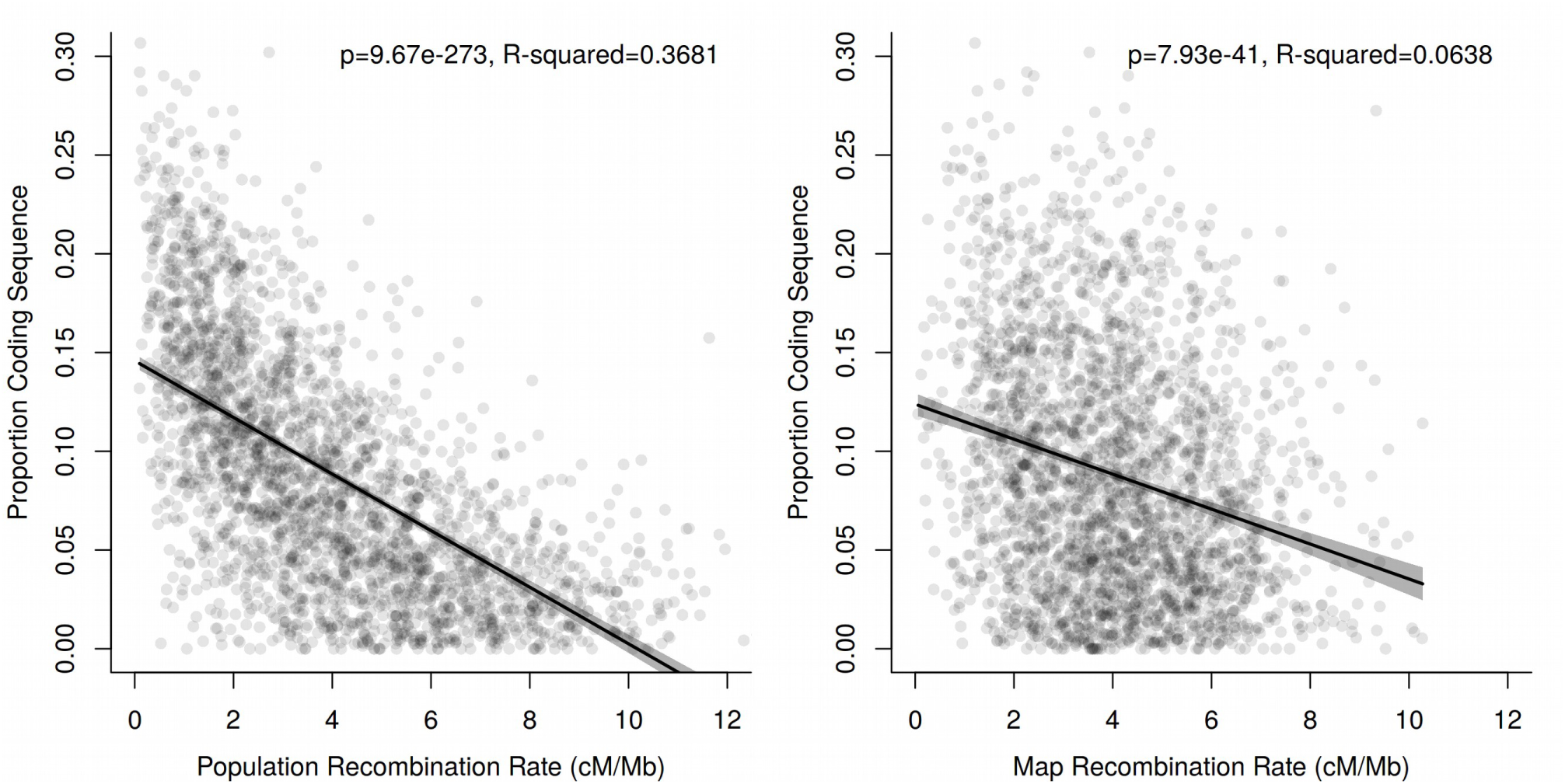
Relationship between recombination rate and gene density. Gene density (i.e. the proportion of coding sequence) in 100 Kb windows plotted against the population recombination rate (ρ) (left) and crossover recombination rate (right). The line shows a fitted linear regression. While there is a strong negative relationship between ρ and gene density, this may partly reflect the fact that ρ represents a composite of recombination and local effective population size, which will tend to be lower in regions of high gene density, due to linked selection [45]. Nevertheless, there is also a negative relationship between the crossover recombination rate and gene density, indicating that regions of lower recombination do indeed tend to harbour more coding sequence. The relationship is fairly weak, but it is unclear to what extent this might reflect the inaccuracies of measuring local recombination rates based on linkage mapping [43].

**Figure S11.**
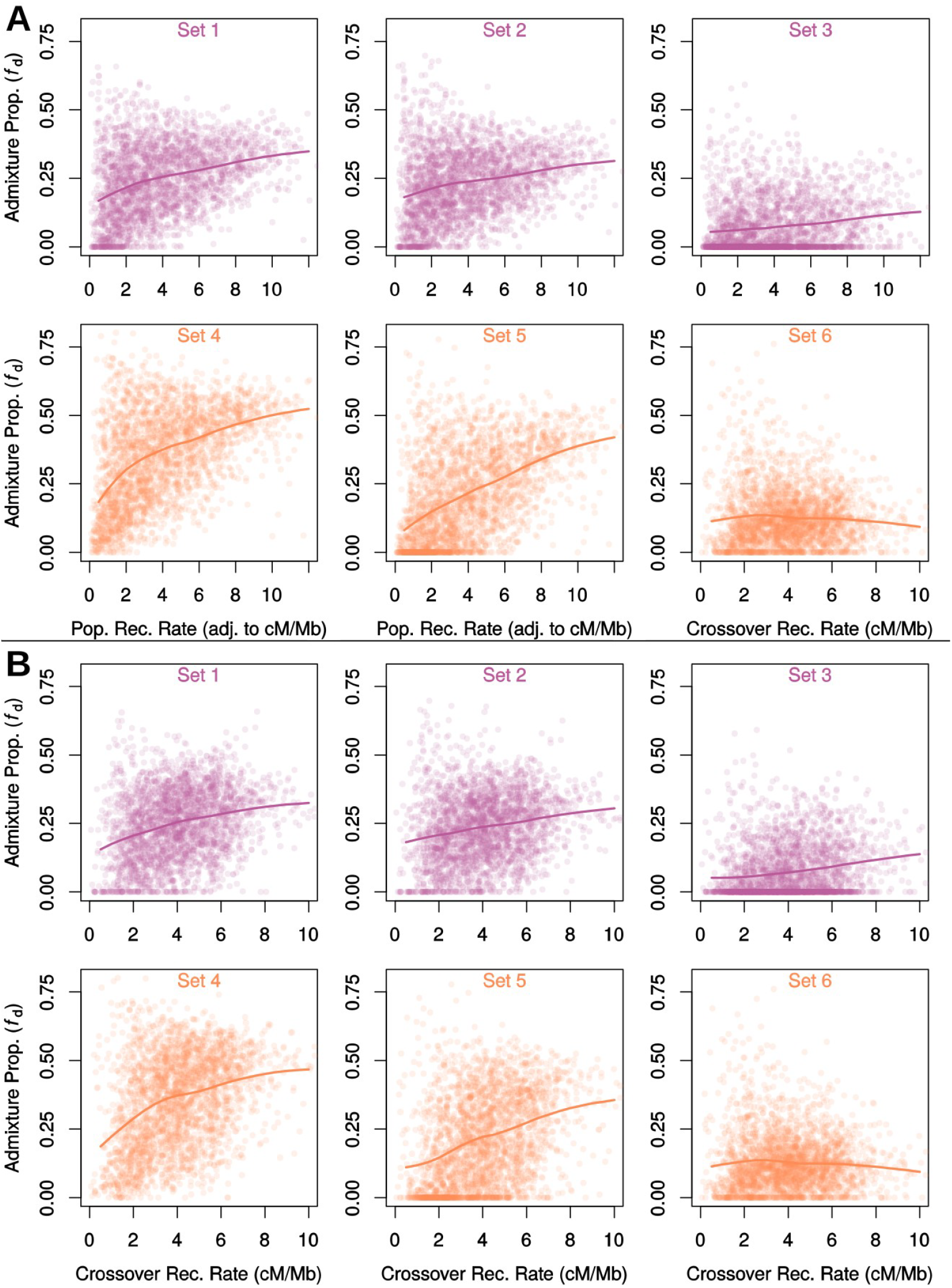
Admixture is positively correlated with recombination rate. Admixture proportions estimated for non-overlapping 100 Kb windows, plotted against the local recombination rate, either computed as the population recombination rate and rescaled to cM/Mb (**A**) or estimated directly from linkage maps (**B**). Solid lines indicate the locally-weighted average (loess span = 0.75). Dashed lines indicate the same average when windows in the outer third of each chromosome are excluded. Admixture between *cyd* and *mel-W* (Sets 1-3, see Figure S7) as well as that between *tim* and *mel-E* (Sets 4-6) increases non-linearly with increasing recombination rate, with the exception of Set 6, for which admixture proportions are low, and there is only evidence for a weak positive relationship in windows of low recombination rate. This may be driven by the close relationship and likely ongoing migration between *mel-E* and *mel-G* (see Figures 1 and S7), which could limit our ability to detect admixture between *tim* and *mel-E*. The estimated admixture proportion between *cyd* and *mel-W* using Set 3 is also much lower than for sets 1 and 2. This may be driven by strongly direction introgression from *cyd* into *mel-W*, which is also indicated by topology weightings, as described in the main paper. If introgression is largely in the direction from P2 into P3, *f*_d_ tends to underestimate the trough admixture proportion [15].

**Figure S12.**
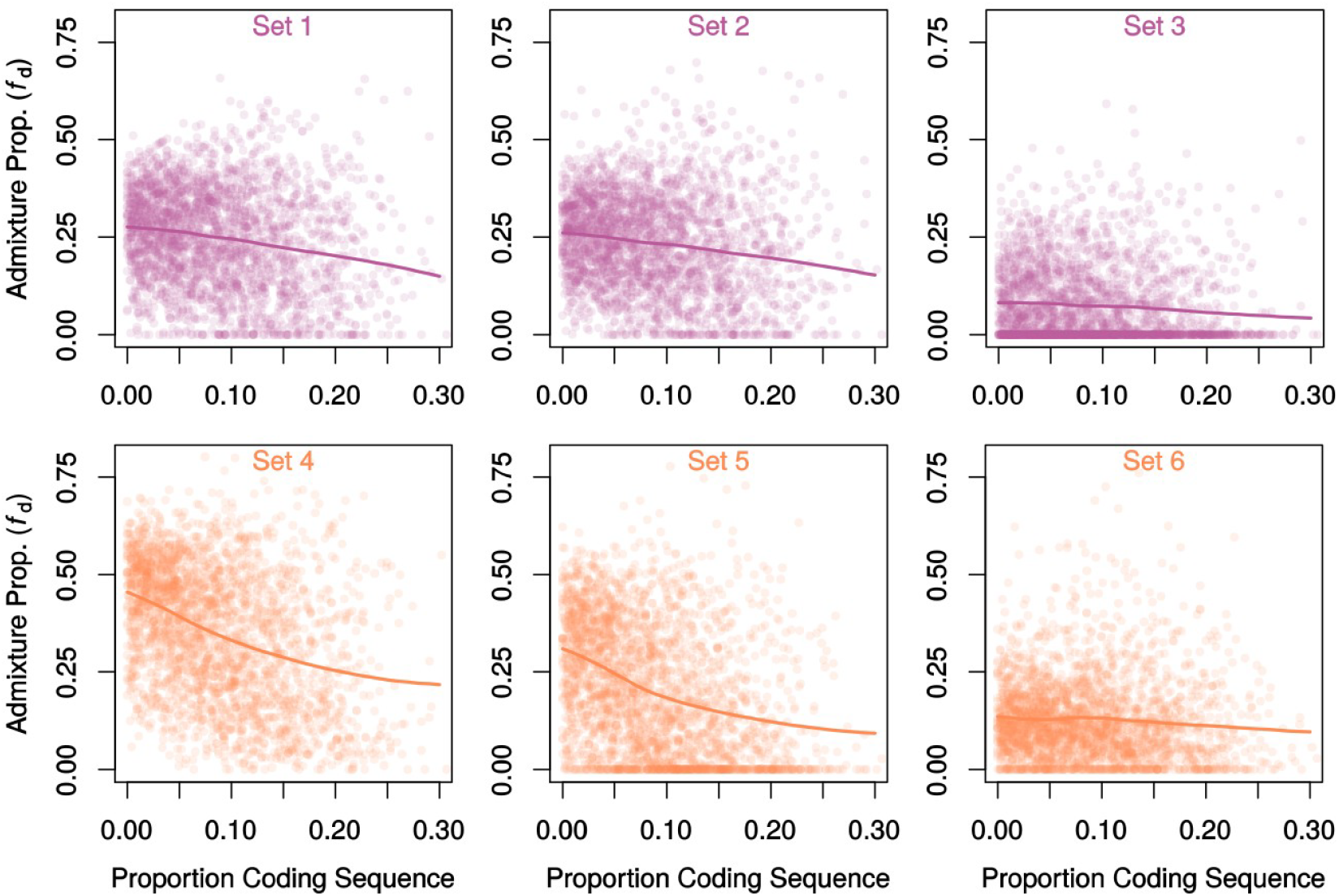
Admixture is negatively correlated with the proportion of coding sequence. Admixture proportions estimated for non-overlapping 100 Kb windows, plotted against the proportion of coding sequence. Solid lines indicate the locally-weighted average (loess span = 0.75). Explanations for the lower average levels of admixture in Sets 3 and 6 are discussed in the legend of Figure S11 above.

